# Physical exercise and motor learning: A scoping review

**DOI:** 10.1101/2025.07.25.666852

**Authors:** Layale Youssef, Amanda O’Farrell, Denis Arvisais, Jason L. Neva, Benjamin Pageaux

## Abstract

Physical exercise can enhance motor learning by inducing neurophysiological changes that facilitate this process. This preregistered scoping review aimed to map current knowledge on the effects of physical exercise on motor learning. Experimental studies and review articles examining any form of physical exercise in healthy or clinical populations were included, with outcomes assessing motor skill acquisition and/or retention. A systematic search across eight databases (e.g., MEDLINE, EMBASE, PsychINFO) identified 66 sources, including 62 experimental studies and 4 reviews. Two researchers independently reviewed articles and extracted data using a pretested data extraction table; disagreements at the study selection or data extraction stages were resolved through discussion with a third researcher. Most studies involved healthy populations (83%), with clinical populations underrepresented (17%). Aerobic exercise was most commonly investigated, particularly lower-limb cycling (73%), while resistance exercise was rarely examined (0.6%). Exercise intensity was predominantly high, although 6% of studies reported intensity not reflecting the prescribed method. Motor learning outcomes varied: 55% assessed both skill acquisition and retention, while 45% relied on retention tests alone at short-term or delayed time points, or both. Infrequent use of long-term retention tests limits understanding of lasting effects. Overall, this review highlights gaps in the literature, including the underrepresentation of clinical populations, inconsistent reporting of exercise intensity, scarce research on resistance exercise, and limited assessment of long-term retention, which may affect interpretation of exercise’s impact on motor learning.

**Highlights:**

- Clinical populations were underrepresented in the exercise and motor learning field.
- High-intensity lower-limb cycling was the most common exercise studied.
- Resistance exercise was underrepresented in the field.
- Accurate prescription of exercise intensity is essential to ensure reliable results.
- Future research should more frequently use longer-term retention tests.

## 1. Introduction

Although the terms “physical activity” and “physical exercise” are sometimes used in similar contexts, they refer to different types of physical movements (Caspersen et al., 1985). Physical activity is defined as any body movement produced by skeletal muscles resulting in energy expenditure (Ndahimana & Kim, 2017). In contrast, physical exercise consists of performing planned and structured physical activity to enhance physical fitness or maintain overall health, and can be either chronic or acute (Kenney et al., 2022). There are different types of physical exercise, including aerobic and resistance exercise. Aerobic exercise includes activities such as running, cycling, and swimming, whereas resistance exercise includes weightlifting and bodyweight exercises. These exercise types can involve different modes of muscular contraction, namely concentric, eccentric, and isometric contractions. Aerobic exercise primarily targets the cardiovascular system and metabolic health, leading to improvement in endurance performance. Resistance exercise primarily targets the neuromuscular system and is essential for developing muscular strength and power (ACSM, 2018). Both aerobic and resistance exercise are essential for maintaining independence in activities of daily living, such as walking, climbing stairs, and carrying objects. Moreover, physical exercise intensity may vary (e.g., low, moderate, and high), and the methodology used to prescribe exercise intensity is different among studies. Importantly, physical exercise is not only beneficial for maintaining physical function and overall health, as it has also been shown to exert beneficial effects on brain health (Cotman & Berchtold, 2002; Phillips et al., 2014; Silva et al., 2024).

Numerous studies in the field of exercise science have highlighted how external interventions, such as physical exercise, can enhance nervous system function (Hillman et al., 2008; Tari et al., 2025). More specifically, physical exercise has been shown to induce significant improvements in cognitive function (Erickson et al., 2011; Phillips et al., 2014). Furthermore, physical exercise has been associated with changes in neuroplasticity (Andrews et al., 2020; Neva, 2025; Neva et al., 2017; Singh et al., 2014; Youssef et al., 2024). Neuroplasticity changes, including alterations in motor cortex excitability, occur during motor learning (Cirillo et al., 2011; Coxon et al., 2014; Holland et al., 2015). Motor cortex excitability reflects the responsiveness, or ease, with which neurons in the motor cortex generate action potentials and trigger muscle activity in response to external stimulation, and is influenced by the balance between excitatory and inhibitory signals that contribute to the overall excitability of the motor cortex (Youssef et al., 2024). Changes in motor cortex excitability suggest that physical exercise may act as a priming mechanism for motor learning, potentially by enhancing motor cortex responsiveness and thereby facilitating the acquisition of new motor skills. Consequently, incorporating physical exercise as an adjunct intervention could be an effective strategy to enhance motor learning (Taubert et al., 2015; Wanner et al., 2020).

A motor skill refers to the ability to perform a goal-directed movement that enables an individual to achieve a specific outcome with accuracy and efficiency, using minimal time and effort (Knapp, 1963). Although often used interchangeably, the terms motor performance and motor learning represent distinct concepts. Motor performance refers to the observable execution of a voluntary action, whereas motor learning involves practice-driven processes that lead to relatively lasting improvements in movement ability (Schmidt & Lee, 2011). Motor learning consists of two fundamental aspects: skill acquisition and skill maintenance. Skill acquisition involves developing movement control through practice, and skill maintenance ensures long-term skill retention and adaptability of the learned skill under different contexts (Krakauer et al., 2019). To study these processes, several motor learning categories have been established, including sequence learning, motor adaptation, motor acuity, and associative learning (for more details, see **Figure S1** in supplementary material 1).

Despite the well-established benefits of physical exercise on physical function and cognition (McMorris et al., 2009; Phillips et al., 2014; Yanagisawa et al., 2010), its impact on motor learning remains unclear, with divergent findings contributing to this uncertainty. For example, some studies found that physical exercise improved motor skill acquisition only (Quaney et al., 2009; Statton et al., 2015), whereas other studies showed improvements in both motor acquisition and motor skill retention following physical exercise (Lehmann et al., 2020; Mang et al., 2014; Neva et al., 2019). Another group of studies reported enhancements only in motor skill retention following physical exercise (Ferrer-Uris et al., 2017; Jespersen et al., 2023; Mang et al., 2016; Stavrinos & Coxon, 2017). In contrast, some studies have found no effect on both skill acquisition and retention following an acute bout of aerobic exercise (Baird et al., 2018; Charalambous et al., 2019; James & Wang, 2023; Pixa et al., 2021; Quinlan et al., 2021; Wanner et al., 2020a). Interestingly, the literature includes a wide range of exercise modalities (e.g., continuous and interval training), types (e.g., aerobic and resistance), and intensities (e.g., low, moderate, and high), as well as varied delays in timing of retention tests (i.e., ranging from hours to days), all of which may contribute to the divergent effects of physical exercise on motor learning. Also, within aerobic exercise, multiple forms have been used, including lower-limb cycling, running, walking, and upper-limb cycling. This diversity in exercise types and protocols adds to the variability observed across studies and underscores the need for careful comparison when interpreting results.

Considering these discrepancies, mapping the existing literature is crucial to better understand the effect of physical exercise on motor learning. A scoping review is particularly suited for this purpose, as it allows researchers to organize and summarize the breadth of evidence, identify key concepts, and uncover gaps in current research, which can then inform future systematic reviews (Munn et al., 2018)

Understanding how different types and intensities of physical exercise impact motor learning is essential for developing targeted exercise protocols that address the specific needs of various populations, whether clinical populations in rehabilitation settings or healthy individuals in sport. Because these populations have distinct characteristics and requirements, tailored exercise protocols may be necessary to optimize motor learning outcomes.

This scoping review aims to explore and map the existing literature on the impact of physical exercise on motor learning in humans. Specifically, we addressed the following research questions:

1. What types of populations were investigated?
2. What types of physical exercise were performed to impact motor learning?
3. When was the physical exercise performed relative to motor task practice?
4. Which types of motor learning categories were used?

These research questions provide a clear structure for understanding the current state of the evidence and highlights areas where further research is needed.

## 2. Methods

This scoping review was i) pre-registered on the Open Science Framework (OSF) Website (https://osf.io/4mv53), ii) conducted according to the JBI scoping review guidelines and recommendations (Peters et al., 2020), and iii) reported according to the PRISMA extension for scoping reviews [PRISMA-ScR, (Tricco et al., 2018)]. The PRISMA-ScR checklist is provided as Supplementary Material 2.

### 2.1 Search strategy

The database search was conducted by a librarian (DA) in collaboration with LY and consultation with BP and JN. The search strategy included a combination of keywords and terms indexed across various databases, including MEDLINE (Ovid), EMBASE (Ovid), PsycINFO (Ovid), SportDiscus with Full Text (Ebsco), CINAHL Complete (Ebsco), ERIC (ProQuest), Web of Science (Clarivate), and Dissertations & Theses Global (ProQuest). The complete search equations are provided in supplementary material 3. The search process was conducted in February 2024 and updated in March 2025. The second search led to the inclusion of two additional experimental studies. The search was limited to articles in English and French, with no restrictions on the publication year. All identified sources were uploaded in Endnote (Clarivate Analytics, PA, USA) and duplicates were removed. The screening of all sources was then managed using Covidence software (Innovation, 2023) and followed three steps: (1) remaining duplicates removal, (2) title and abstract screening, and (3) full-text evaluation. Two researchers (LY and AOF) independently reviewed articles at each stage, and any disagreements were resolved through discussion with a third researcher (BP). A detailed flowchart outlining the selection process of sources can be found in **Figure 1**.

**Figure 1.**
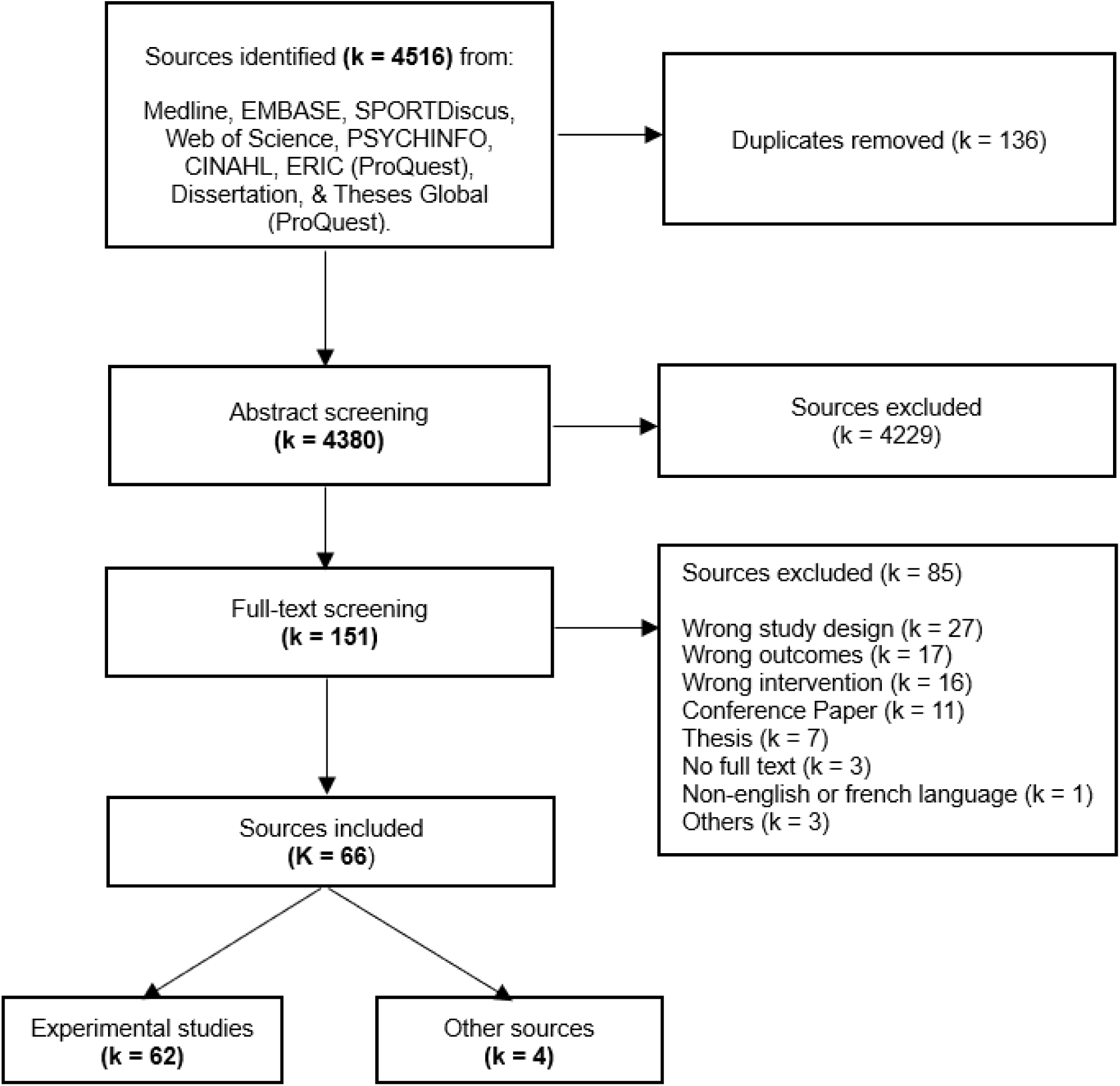
Flowchart illustrating the selection process of sources.

### 2.2 Inclusion Criteria

All fully published, peer-reviewed articles evaluating the effect of physical exercise on motor learning were included. Both experimental studies and reviews that incorporated experimental studies were considered. The primary focus was the impact of physical exercise on motor learning, assessed in either laboratory or field settings. Studies were included if they involved clearly defined motor tasks with assessments of skill acquisition, with or without retention tests. However, following the classical definition of motor learning (Schmidt & Lee, 2011), skill maintenance over time was not considered in studies where a retention test was not included. Furthermore, all types of physical exercise interventions (aerobic or resistance) at any intensity level (low, moderate, high) were included. Finally, the review included studies involving all types of human populations (healthy and clinical) of all ages.

### 2.3 Data extraction

The extracted data included details about the evaluated populations, the physical exercise interventions, the motor learning parameters and additional key findings relevant to the objectives of this scoping review. Data extraction was carried out by two independent reviewers (LY and AOF), with any disagreements resolved through discussion. If consensus could not be reached, JN and BP were consulted to provide a final resolution. A data extraction table was used to record the information from the included studies. The initial table was tested on two articles by LY and AOF, then slightly refined before reaching its final version presented in supplementary material 4. For experimental studies, the following data were collected: population characteristics (age and type), details of the physical exercise performed (type, duration, intensity) based on American College of Sports Medicine (ACSM) guidelines (ACSM, 2018), timing of exercise relative to motor task practice (before or after acquisition), motor learning category (sequence, motor acuity, motor adaptation, associative learning) as well as the result as reported in each study (i.e, effect of physical exercise on skill acquisition and/or skill retention). For review articles, data extraction was limited to qualitative information only, specifically the main conclusions and take-home messages related to the effects of physical exercise on motor learning. No quantitative data were extracted from review papers, and primary data reported within these reviews were not re-extracted, thereby avoiding duplication or redundancy with the data extracted in the experimental studies.

Although we followed the pre-registration for the review, two minor deviations were made during the data extraction process to improve methodological clarity. First, we introduced an additional distinction between studies regarding the methodology for exercise prescription. We decided to dissociate prescriptions based on measured versus estimated parameters (e.g., estimated vs measured maximal heart rate). This refinement was not initially planned but was added during data extraction to better capture variability in exercise prescription methods across studies. Second, regarding the motor learning categories (Krakauer et al., 2019), we originally intended to use predefined categories such as sequence learning and adaptation learning. However, after examining the motor tasks used in the included studies, we found that these predefined categories did not fully capture the distinct characteristics of the motor learning constructs investigated. Therefore, we refined our categorization to better reflect the outcomes being evaluated. The categorization used is detailed in **Figure S1** of supplementary material 1.

In this scoping review, the number of experimental conditions exceeds the number of studies, as several studies included multiple experimental conditions. For example, some studies involved different exercise intensities or different populations. As a result, the mapping of the literature was also conducted based on the number of experimental conditions relevant to each research question. In the manuscript, *n* refers to a subtotal of the number of participants, *k* refers to a subtotal of the number of studies, *g* refers to a subtotal of the number of groups, and *e* refers to a subtotal of the number of experimental conditions. When these letters are capitalized, it refers to the grand total (e.g., N = the total number of participants including all studies and all experimental conditions).

### Motor learning categorization and results

To classify the motor learning categories used across studies, we identified four main categories based on task characteristics and learning objectives (Krakauer et al., 2019). *Sequence learning*, often referred to as sequence-specific learning, was defined for motor tasks that involve comparing performance on a repeated sequence or sequences with that of random sequences (e.g., serial reaction time task). This comparison is necessary to identify learning that is specific to the practiced sequence rather than general improvements in other aspects of the motor skill. However, if only a repeated sequence or sequences were practiced without including random sequences for comparison, the motor task was categorized as motor acuity. *Motor acuity* refers to improvements in the accuracy and/or precision of performing a particular movement or sequence of movements through practice (e.g., Rho task). In this case, the focus is on refining the ability to execute the motor task more precisely and/or accurately. *Motor adaptation* included tasks in which participants adjusted their movements in response to external perturbation (e.g., visuomotor rotation task). Finally, *associative learning* referred to tasks requiring participants to form arbitrary associations between stimuli and responses (e.g., conditional learning task). The primary objective of each category is shown in **Figure S1** of supplementary material This classification provides a clear view of the distribution of motor learning categories across the included studies and the identification of gaps in literature.

To provide a comprehensive understanding of the effect of physical exercise on motor learning, we examined skill acquisition and skill retention separately based on when physical exercise was performed relative to the motor task. The results for both skill acquisition and skill retention were reported in studies that performed physical exercise prior to the motor task, while only results from retention tests were reported in studies that conducted physical exercise after the motor task. Furthermore, the timing of retention tests varied across studies. To address this potential challenge in the synthesis of findings, we categorized the reported results from retention tests into three time windows: before 24 hours, at 24 hours, and after 24 hours. Consequently, findings from some studies appear in multiple retention time windows. Results were categorized based on exercise intensity and separated ccording to population type. However, due to the complexity of intervention types and timing, all forms of physical exercise were grouped together in the outcome of this scoping review, regardless of when the exercise was performed. To accurately assess the effects of physical exercise on motor learning, a comparison with a control condition (i.e., without physical exercise) is essential and was included in the majority of the studies. Based on the results reported by the authors in the experimental studies, motor learning effects were categorized as enhanced, no difference, or attenuated performance. Although studies without a control condition were included in the scoping review and their data were extracted, their findings were not considered in the outcome of this scoping review.

## 3. Results

### 3.1 Included studies

Following the systematic search in various databases, 66 sources were included (see **Figure 1**). Following the JBI guidelines for categorization, 62 primary sources were experimental studies and 4 secondary sources were review articles [1 narrative review (Taubert et al., 2015), 2 systematic reviews (Bonuzzi & Torriani-Pasin, 2022; Hubner & Voelcker-Rehage, 2017), and 1 systematic review with meta-analysis (Wanner et al., 2020)]. A timeline of the included sources is presented in **Figure 2a**. The research interest on the effect of physical exercise on motor learning first appeared as early as 1970, but after a long gap, studies in this area began to emerge more consistently starting in 2009, reaching a peak in 2023 with 13 studies published that year. A world map illustrating the geographical distribution of the included studies is presented in **Figure 2b**. This scoping review revealed a worldwide interest in the effects of physical exercise on motor learning, with most studies conducted in Canada (k = 16), the United States (k = 11), Germany (k = 10) and Denmark (k = 6).

**Figure 2.**
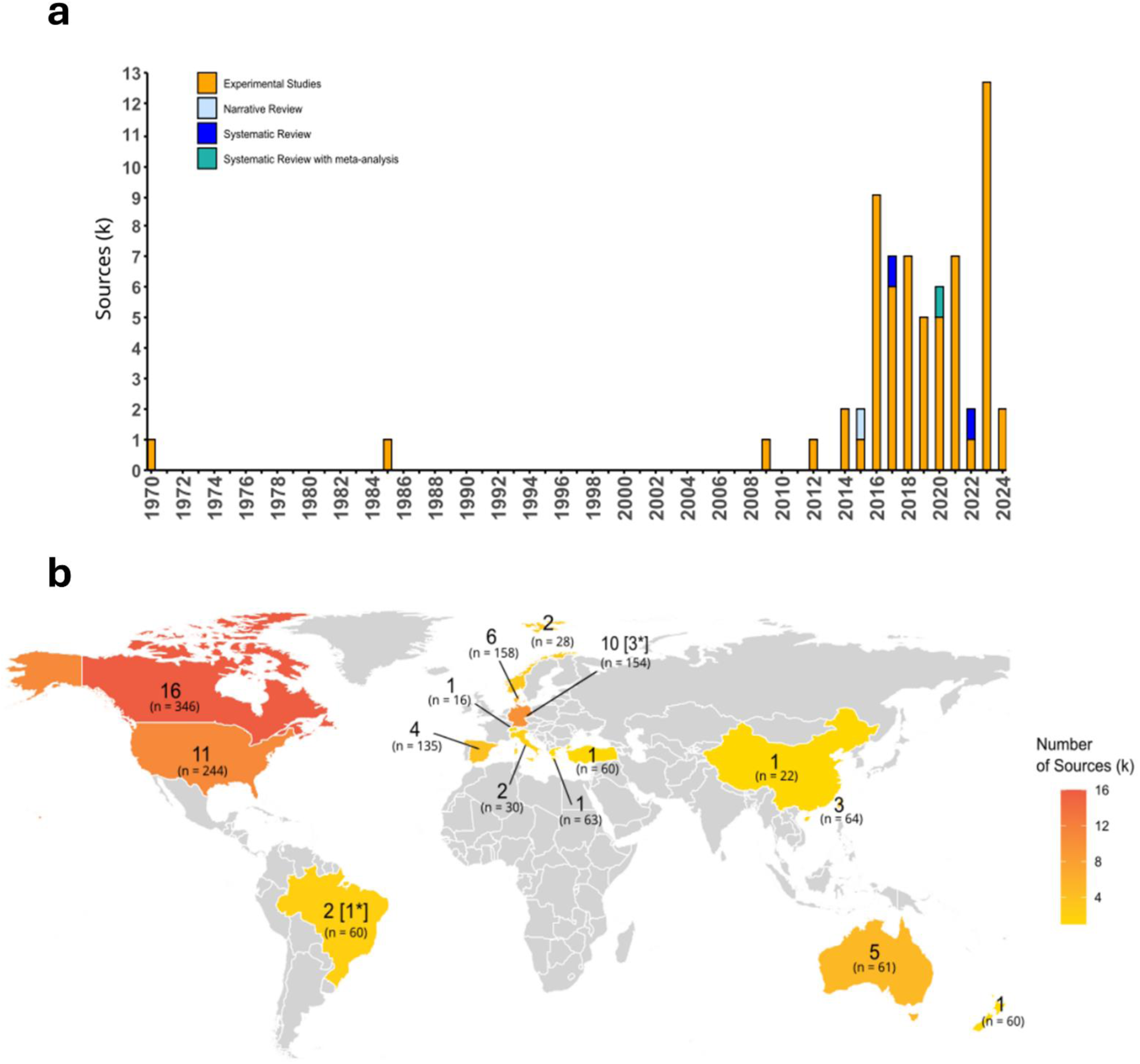
Temporal and geographical distribution of included sources. (Panel a) Temporal distribution of all included sources. Orange bars represent experimental studies, while blue gradient bars represent other types of sources (narrative review; systematic review; systematic review with meta-analysis). (Panel b) Geographical distribution of all included sources. The format used is the following: total number [other types of sources*]. The total number of sources is shown first; when sources other than experimental studies are included, their number is indicated in brackets. Darker colors indicate a higher number of sources reported from each country. n refers to the number of participants reported from experimental studies conducted in that country.

We extracted data from 62 experimental studies to address our specific research questions. The following sections address our review questions, detailing the populations studied, the types of physical exercise implemented, the motor learning parameters investigated, and the timing of physical exercise in relation to motor task practice. An overview of the main characteristics of all experimental studies is presented in **Table 1**. Detailed characteristics of all included experimental studies are provided in **Table S1** (Supplementary Material 1), which offers comprehensive study-level information (e.g., population characteristics, exercise protocols, and outcomes). This table is intended for readers interested in exploring individual studies in greater detail.

**Table 1:**
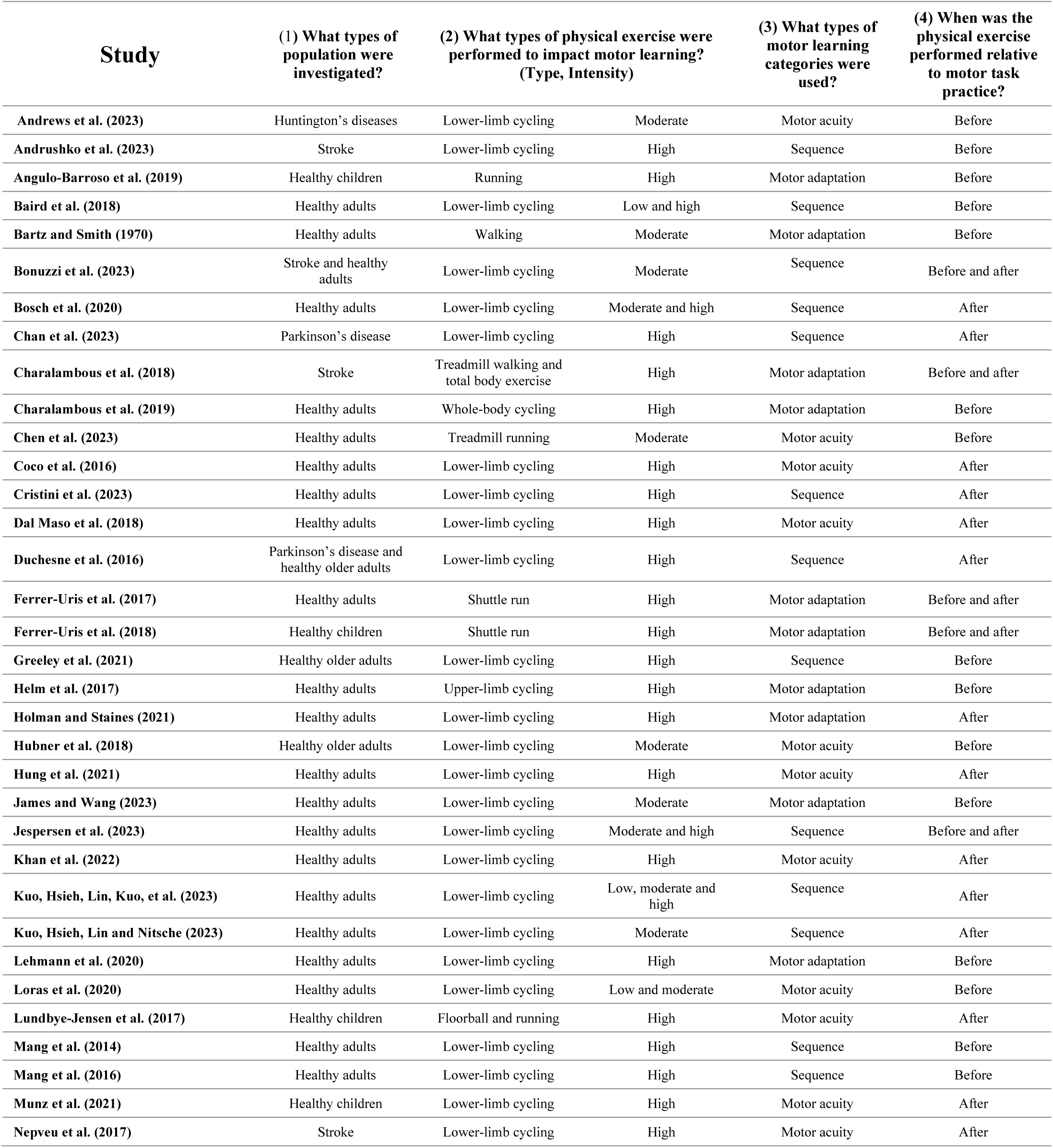
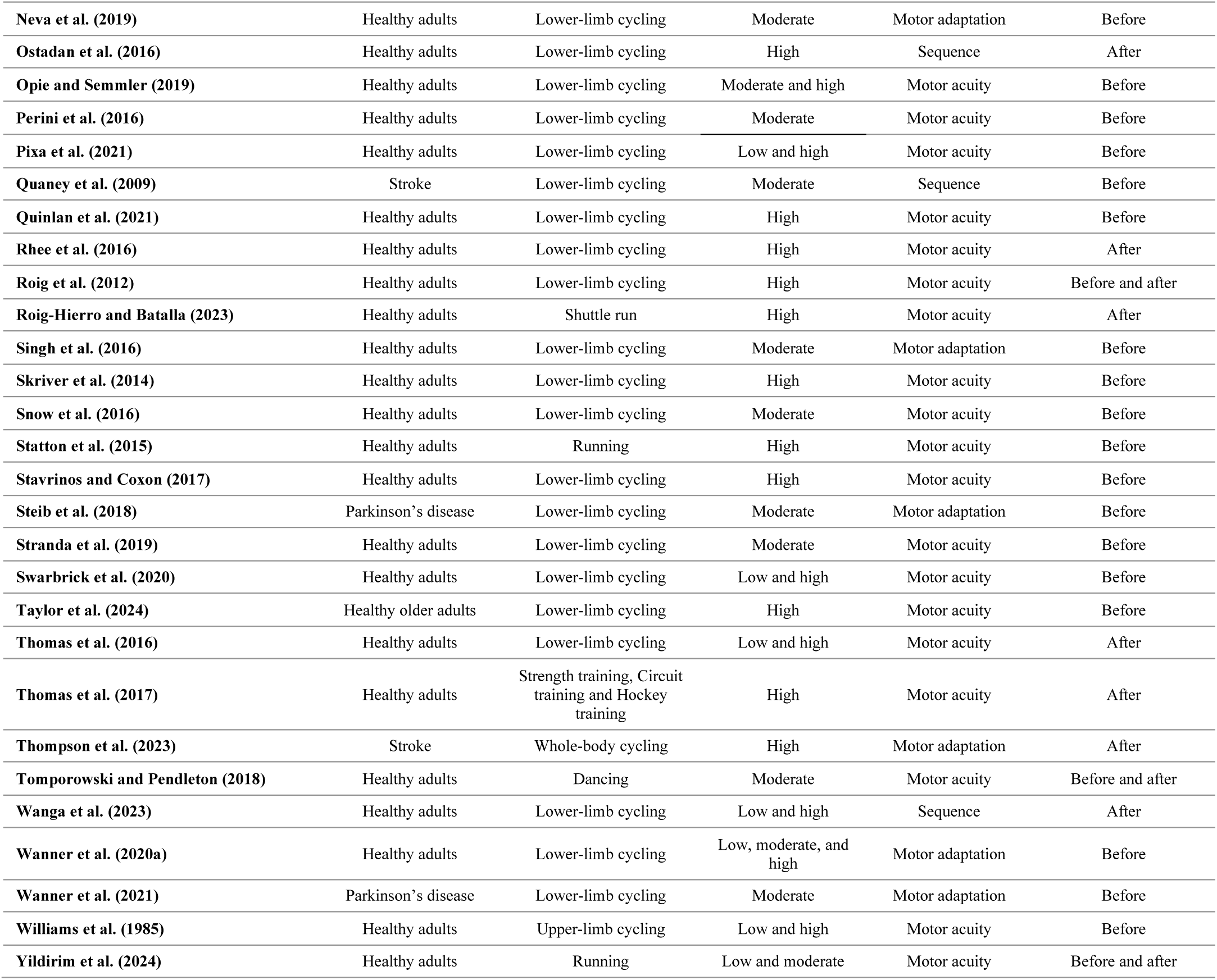
Characteristics of the included studies related to the review questions.

| Study | (1) What types of population were investigated? | (2) What types of physical exercise were performed to impact motor learning? (Type, Intensity) |  | (3) What types of motor learning categories were used? | (4) When was the physical exercise performed relative to motor task practice? |
| --- | --- | --- | --- | --- | --- |
| Andrews et al. (2023) | Huntington’s diseases | Lower-limb cycling | Moderate | Motor acuity | Before |
| Andrushko et al. (2023) | Stroke | Lower-limb cycling | High | Sequence | Before |
| Angulo-Barroso et al. (2019) | Healthy children | Running | High | Motor adaptation | Before |
| Baird et al. (2018) | Healthy adults | Lower-limb cycling | Low and high | Sequence | Before |
| Bartz and Smith (1970) | Healthy adults | Walking | Moderate | Motor adaptation | Before |
| Bonuzzi et al. (2023) | Stroke and healthy adults | Lower-limb cycling | Moderate | Sequence | Before and after |
| Bosch et al. (2020) | Healthy adults | Lower-limb cycling | Moderate and high | Sequence | After |
| Chan et al. (2023) | Parkinson’s disease | Lower-limb cycling | High | Sequence | After |
| Charalambous et al. (2018) | Stroke | Treadmill walking and total body exercise | High | Motor adaptation | Before and after |
| Charalambous et al. (2019) | Healthy adults | Whole-body cycling | High | Motor adaptation | Before |
| Chen et al. (2023) | Healthy adults | Treadmill running | Moderate | Motor acuity | Before |
| Coco et al. (2016) | Healthy adults | Lower-limb cycling | High | Motor acuity | After |
| Cristini et al. (2023) | Healthy adults | Lower-limb cycling | High | Sequence | After |
| Dal Maso et al. (2018) | Healthy adults | Lower-limb cycling | High | Motor acuity | After |
| Duchesne et al. (2016) | Parkinson’s disease and healthy older adults | Lower-limb cycling | High | Sequence | After |
| Ferrer-Uris et al. (2017) | Healthy adults | Shuttle run | High | Motor adaptation | Before and after |
| Ferrer-Uris et al. (2018) | Healthy children | Shuttle run | High | Motor adaptation | Before and after |
| Greeley et al. (2021) | Healthy older adults | Lower-limb cycling | High | Sequence | Before |
| Helm et al. (2017) | Healthy adults | Upper-limb cycling | High | Motor adaptation | Before |
| Holman and Staines (2021) | Healthy adults | Lower-limb cycling | High | Motor adaptation | After |
| Hubner et al. (2018) | Healthy older adults | Lower-limb cycling | Moderate | Motor acuity | Before |
| Hung et al. (2021) | Healthy adults | Lower-limb cycling | High | Motor acuity | After |
| James and Wang (2023) | Healthy adults | Lower-limb cycling | Moderate | Motor adaptation | Before |
| Jespersen et al. (2023) | Healthy adults | Lower-limb cycling | Moderate and high | Sequence | Before and after |
| Khan et al. (2022) | Healthy adults | Lower-limb cycling | High | Motor acuity | After |
| Kuo, Hsieh, Lin, Kuo, et al. (2023) | Healthy adults | Lower-limb cycling | Low, moderate and high | Sequence | After |
| Kuo, Hsieh, Lin and Nitsche (2023) | Healthy adults | Lower-limb cycling | Moderate | Sequence | After |
| Lehmann et al. (2020) | Healthy adults | Lower-limb cycling | High | Motor adaptation | Before |
| Loras et al. (2020) | Healthy adults | Lower-limb cycling | Low and moderate | Motor acuity | Before |
| Lundbye-Jensen et al. (2017) | Healthy children | Floorball and running | High | Motor acuity | After |
| Mang et al. (2014) | Healthy adults | Lower-limb cycling | High | Sequence | Before |
| Mang et al. (2016) | Healthy adults | Lower-limb cycling | High | Sequence | Before |
| Munz et al. (2021) | Healthy children | Lower-limb cycling | High | Motor acuity | After |
| Nepveu et al. (2017) | Stroke | Lower-limb cycling | High | Motor acuity | After |

**Table 1. Continued**
| Study | (1) What types of population were investigated? | (2) What types of physical exercise were performed to impact motor learning? (Type, Intensity) |  | (3) What types of motor learning categories were used? | (4) When was the physical exercise performed relative to motor task practice? |
| --- | --- | --- | --- | --- | --- |
| Neva et al. (2019) | Healthy adults | Lower-limb cycling | Moderate | Motor adaptation | Before |
| Ostadan et al. (2016) | Healthy adults | Lower-limb cycling | High | Sequence | After |
| Opie and Semmler (2019) | Healthy adults | Lower-limb cycling | Moderate and high | Motor acuity | Before |
| Perini et al. (2016) | Healthy adults | Lower-limb cycling | Moderate | Motor acuity | Before |
| Pixa et al. (2021) | Healthy adults | Lower-limb cycling | Low and high | Motor acuity | Before |
| Quaney et al. (2009) | Stroke | Lower-limb cycling | Moderate | Sequence | Before |
| Quinlan et al. (2021) | Healthy adults | Lower-limb cycling | High | Motor acuity | Before |
| Rhee et al. (2016) | Healthy adults | Lower-limb cycling | High | Motor acuity | After |
| Roig et al. (2012) | Healthy adults | Lower-limb cycling | High | Motor acuity | Before and after |
| Roig-Hierro and Batalla (2023) | Healthy adults | Shuttle run | High | Motor acuity | After |
| Singh et al. (2016) | Healthy adults | Lower-limb cycling | Moderate | Motor adaptation | Before |
| Skriver et al. (2014) | Healthy adults | Lower-limb cycling | High | Motor acuity | Before |
| Snow et al. (2016) | Healthy adults | Lower-limb cycling | Moderate | Motor acuity | Before |
| Statton et al. (2015) | Healthy adults | Running | High | Motor acuity | Before |
| Stavrinos and Coxon (2017) | Healthy adults | Lower-limb cycling | High | Motor acuity | Before |
| Steib et al. (2018) | Parkinson's disease | Lower-limb cycling | Moderate | Motor adaptation | Before |
| Stranda et al. (2019) | Healthy adults | Lower-limb cycling | Moderate | Motor acuity | Before |
| Swarbrick et al. (2020) | Healthy adults | Lower-limb cycling | Low and high | Motor acuity | Before |
| Taylor et al. (2024) | Healthy older adults | Lower-limb cycling | High | Motor acuity | Before |
| Thomas et al. (2016) | Healthy adults | Lower-limb cycling | Low and high | Motor acuity | After |
| Thomas et al. (2017) | Healthy adults | Strength training, Circuit training and Hockey training | High | Motor acuity | After |
| Thompson et al. (2023) | Stroke | Whole-body cycling | High | Motor adaptation | After |
| Tompowski and Pendleton (2018) | Healthy adults | Dancing | Moderate | Motor acuity | Before and after |
| Wanga et al. (2023) | Healthy adults | Lower-limb cycling | Low and high | Sequence | After |
| Wanner et al. (2020a) | Healthy adults | Lower-limb cycling | Low, moderate, and high | Motor adaptation | Before |
| Wanner et al. (2021) | Parkinson's disease | Lower-limb cycling | Moderate | Motor adaptation | Before |
| Williams et al. (1985) | Healthy adults | Upper-limb cycling | Low and high | Motor acuity | Before |
| Yildirim et al. (2024) | Healthy adults | Running | Low and moderate | Motor acuity | Before and after |

### 3.2 What types of populations were investigated?

The majority of included studies investigated healthy populations (g = 53), while only a small portion of studies (g = 11) focused on clinical populations (see **Figure 3**). Among the studies including healthy populations, 45 studies involved young healthy adults (studies are listed in **Table 1**). Research involving other age groups was limited, with only 4 studies investigating healthy children (Angulo-Barroso et al., 2019; Ferrer-Uris et al., 2018; Lundbye-Jensen et al., 2017; Munz et al., 2021) and 4 studies focusing on healthy older adults (Duchesne et al., 2015; Greeley et al., 2021; Hubner et al., 2018; Taylor et al., 2024). Within the studies investigating clinical populations, three neurological conditions were identified with 6 studies investigating individuals with stroke (Andrushko et al., 2023; Bonuzzi et al., 2023; Charalambous et al., 2018; Nepveu et al., 2017; Quaney et al., 2009; Thompson et al., 2023), followed by 4 studies investigating individuals with Parkinson’s disease (Chan et al., 2023; Duchesne et al., 2015; Steib et al., 2018; Wanner et al., 2021). Additionally, a single study focused on individuals with Huntington’s disease (Andrews et al., 2023).

**Figure 3.**
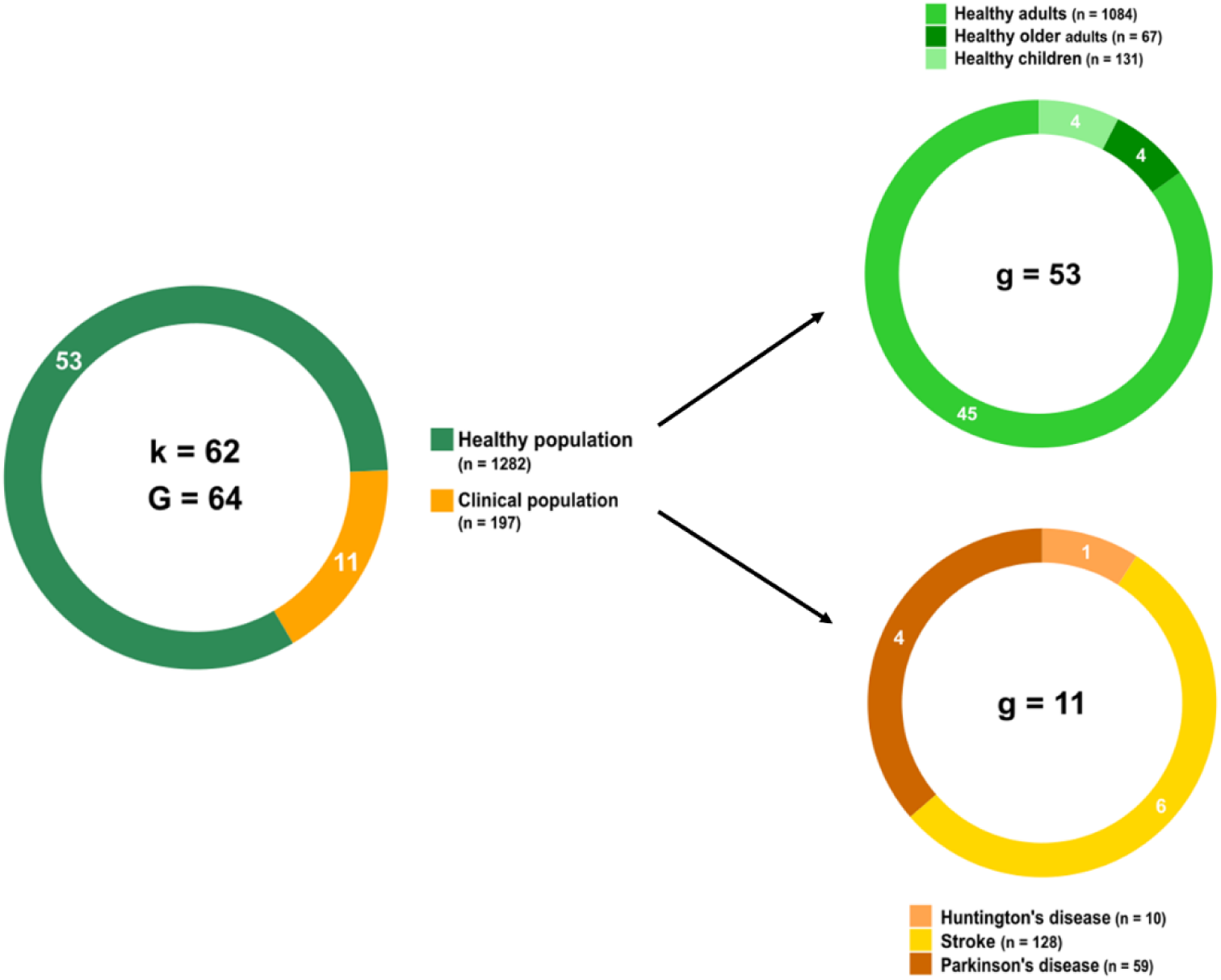
Types of populations evaluated across all included experimental studies. Distribution of the 1,479 participants reported across the 62 experimental studies. Participants were categorized as either healthy or clinical populations, with each category further divided into subcategories. Green gradients represent healthy populations, and orange gradients represent clinical populations. k refers to the number of experimental studies, G refers to the overall number of reported groups, g refers to the number of reported subgroups.

### 3.3 What types and intensities of physical exercise were used?

Among the 62 experimental studies, a total of 81 experimental conditions were conducted (E = 81). The vast majority involved aerobic exercise (e = 79), while only a small number focused on resistance exercise (e = 2) (see **Figure 4a**). Of the aerobic exercise conditions, lower-limb cycling was the main type of exercise used (e = 59). Other aerobic exercise modalities included running (Angulo-Barroso et al., 2019; Chen et al., 2023; Ferrer-Uris et al., 2018; Ferrer-Uris et al., 2017; Lundbye-Jensen et al., 2017; Roig-Hierro & Batalla, 2023; Statton et al., 2015; Yildirim et al., 2024), walking (Bartz & Smith, 1970; Charalambous et al., 2018), upper-limb cycling (Helm et al., 2017; Williams et al., 1985), combined upper and lower-limb cycling (whole-body cycling) (Charalambous et al., 2019; Thompson et al., 2023), and whole-body aerobic exercise (Charalambous et al., 2018). Additionally, a small number of studies employed less common physical exercise, including floorball (Lundbye-Jensen et al., 2017), dancing (Tomporowski & Pendleton, 2018), and hockey (Thomas et al., 2017).

**Figure 4.**
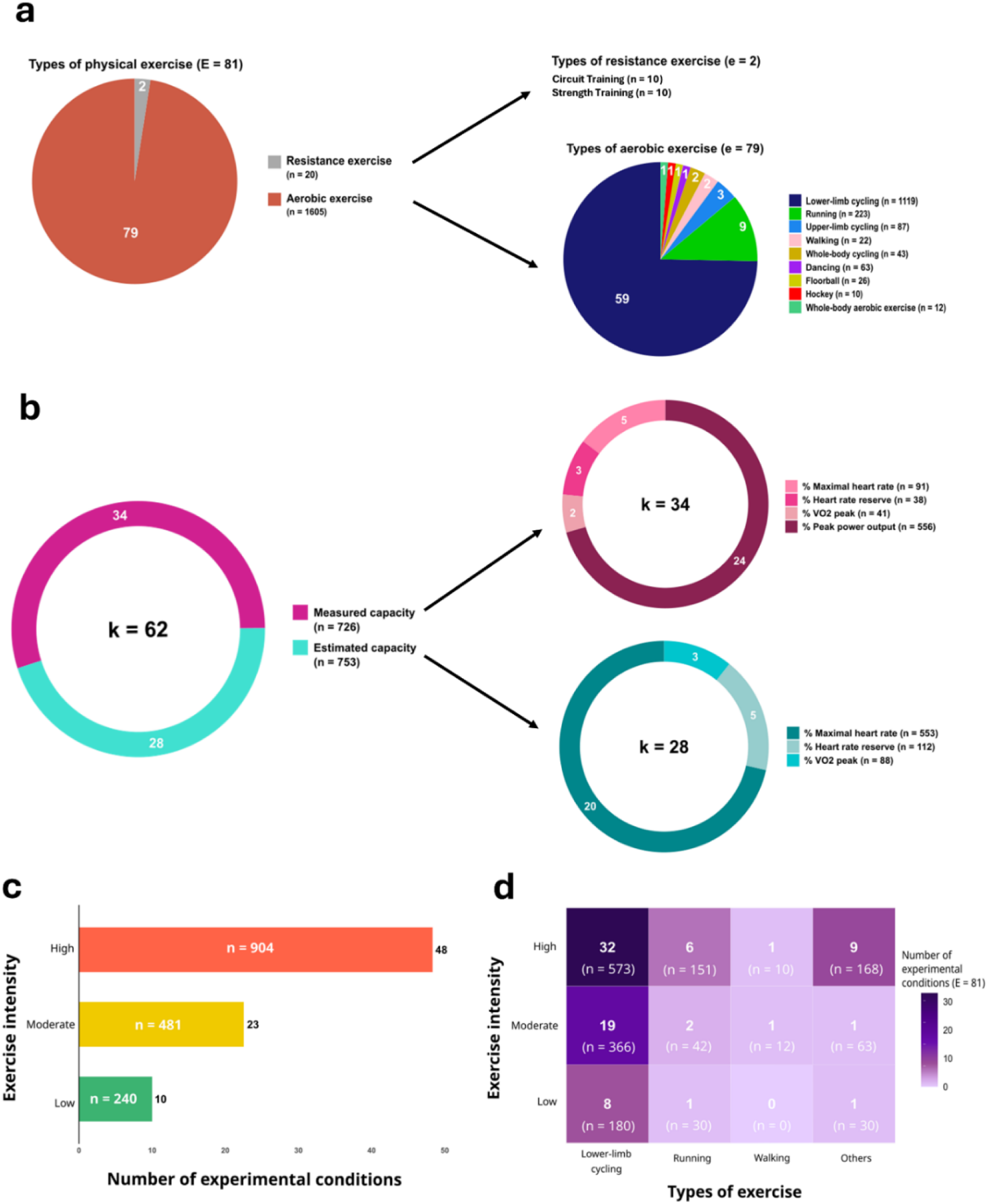
Types and intensities of physical exercise reported across all included experimental studies. (Panel a) Distribution of the 81 experimental conditions according to exercise type, categorized as aerobic or resistance. Aerobic exercise is further subcategorized by specific types. (Panel b) Prescription of exercise intensities across the experimental studies, pink refers to measured capacity and turquoise refers to estimated capacity. Pink and turquoise gradients refer to measured and estimated exersie parameters respectively. (Panel c) Exercise intensities used across all experiments where red refers to high intensity, yellow refers to moderate intensity and green refers to low intensity. (Panel d) Heatmap showing the combination of exercise type and intensity where darker purple indicates a higher number of experimental conditions using the specific combination. Others refers to all exercise types different from lower-limb cycling, running and walking. E refers to the total number of experimental conditionseported across the 62 included experimental studies, e refers to the number of experimental conditions per exercise type category and n refers to the number of participants.

Of the 62 experimental studies included, 34 prescribed exercise intensity based on measured physiological parameters reflecting individual capacity while 28 relied on estimated parameters (see **Figure 4b**). No study used the perception of effort or other perceptual responses to prescribe the exercise intensity. According to ACSM guidelines, the 81 experimental conditions were further classified as high intensity (e = 48), moderate intensity (e = 23), and low intensity (e = 10) (see **Figure 4c**). When combining exercise type and intensity, lower-limb cycling at high intensity was the most commonly used combination, followed by lower-limb cycling at moderate intensity (see **Figure 4d**).

Among the 81 experimental conditions, five showed discrepancies between the reported and actual exercise intensities, as defined by the ACSM guidelines. Specifically, two conditions were reported as moderate intensity, but the prescribed workloads—70% age-predicted heart rate reserve (Holman & Staines, 2021) and 65-85% age-predicted maximal heart rate with a reported average of 84% maximal heart rate (Statton et al., 2015)—fall within the high-intensity range. In another study, two conditions were labeled as moderate and high intensity, but the prescriptions of 50% age-predicted maximal heart rate and 75% age-predicted maximal heart rate respectively, actually correspond to low and moderate intensity based on ACSM guidelines (Loras et al., 2020). Also, one condition was described as low intensity, although the exercise was performed at 50% age-predicted heart rate reserve, which is interpreted as moderate intensity (Opie & Semmler, 2019).

### 3.4 Which categories of motor learning were investigated, and when was physical exercise performed relative to motor task practice?

Three motor learning categories were used across the 62 included experimental studies (see **Figure 5a**): 30 involved motor acuity, 16 focused on sequence learning, and 16 addressed motor adaptation; no study investigated associative learning (studies are listed in **Table 1**).

**Figure 5.**
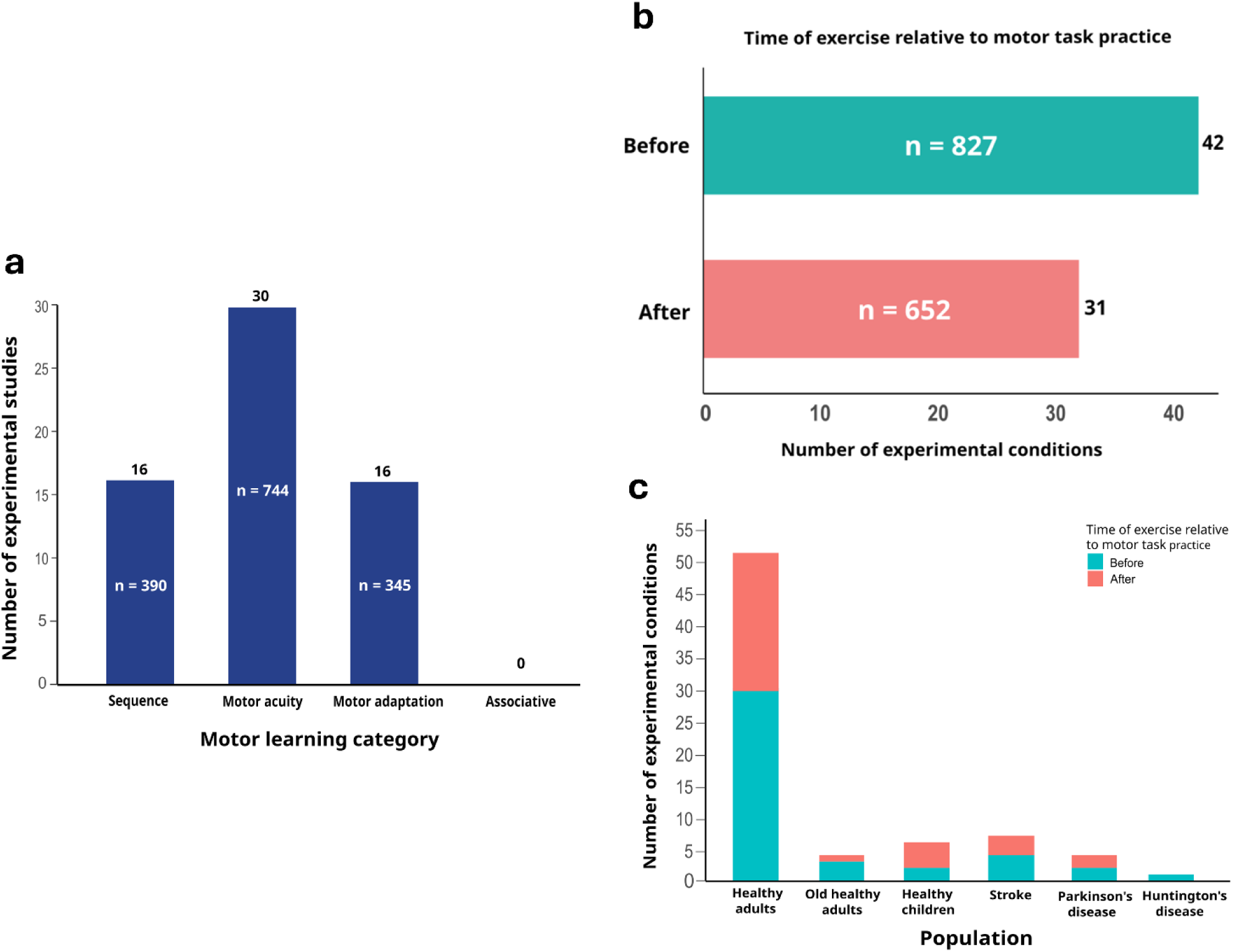
Motor learning categories and timing of exercise relative to motor task practice. (Panel a) Distribution of motor learning categories evaluated across the 62 experimental studies. (Panel b) Timing of exercise in relation to motor task practice across the 73 experimental conditions, (Panel c) Timing of exercise in relation to motor task practice across the 73 experimental conditions shown by population type. Light blue indicates exercise performed before the motor task practice, and light red indicates exercise performed after the motor task practice. Before = exercise performed prior to motor task practice; After = exercise performed after the motor task practice; n refers to the number of participants.

Among the 73 experimental conditions related to the timing of the exercise performed (E = 73), 42 involved physical exercise performed before motor task practice and 31 after (see **Figure 5b**). In healthy adults (e = 51), 30 conditions implemented exercise before motor task practice and 21 after. Among older healthy adults (e = 4), three conditions performed exercise before and one after motor task practice. In healthy children (e = 6), two conditions included exercise before and four after motor task practice. In individuals with stroke (e = 7), four conditions involved exercise before and three after motor task practice. In individuals with Parkinson’s disease (e = 4), two conditions implemented exercise before and two after motor task practice. Finally, the single condition involving individuals with Huntington’s disease (e = 1) performed exercise before motor task practice (see **Figure 5c**).

### 3.5 Effects of physical exercise on motor learning

The effects of physical exercise on motor learning, as reported by the authors in each included experimental study, were categorized based on acquisition and retention tests (enhanced, no difference, or attenuated) relative to a control condition. Results for the retention tests were separated into three time windows: less than 24 hours, at 24 hours, and more than 24 hours. These results were categorized by exercise intensity (high, moderate and low) and by population type (see **Figure 6**). Detailed results for each study are also provided in **Table S1** of supplementary material 1.

**Figure 6.**
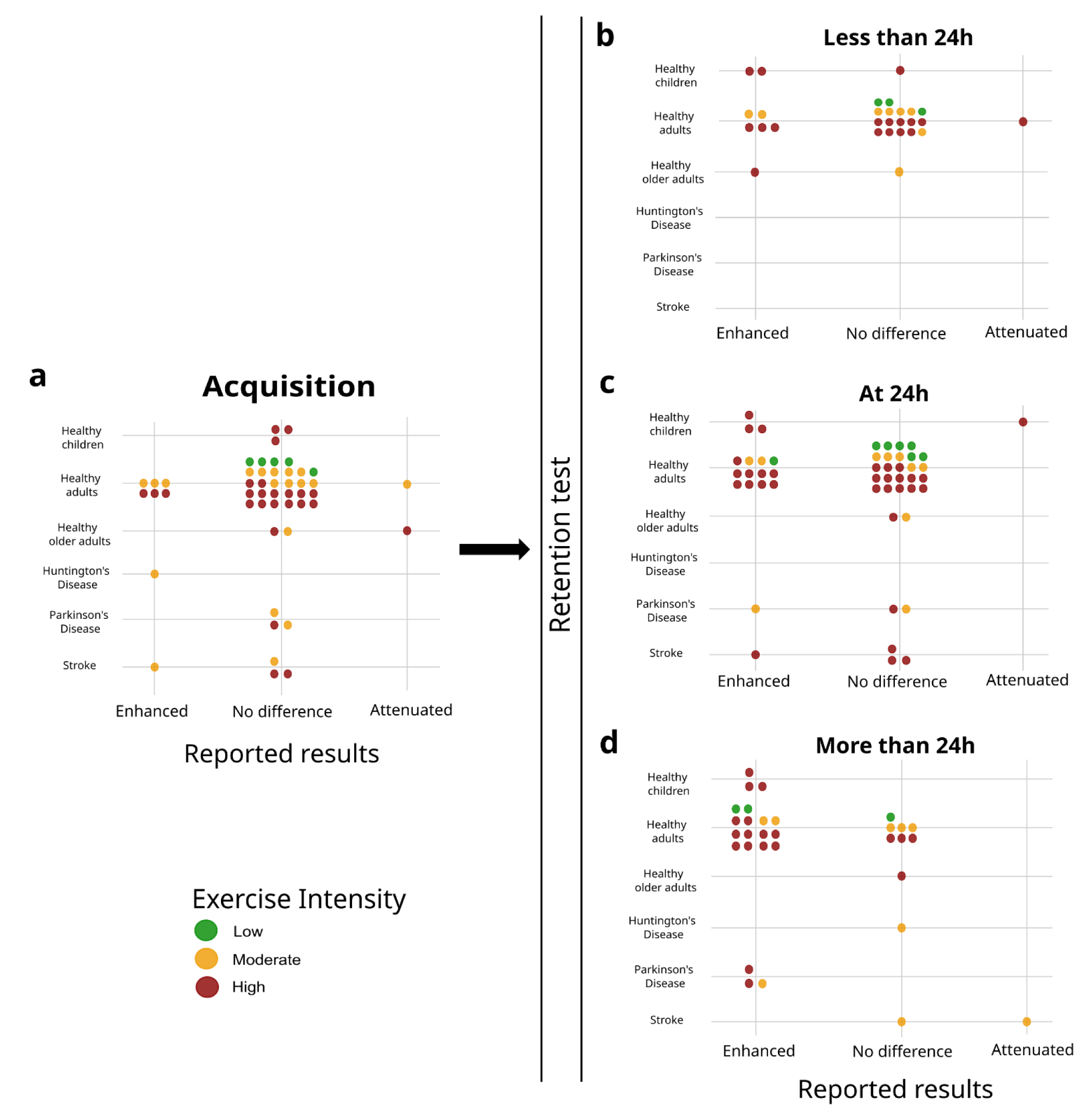
Effects of physical exercise interventions on motor learning. Colored dots represent individual experimental conditions, with their position indicating whether motor learning was enhanced, not different, or attenuated compared to a control condition. Only experiments including a control condition were considered. All population types investigated are represented. Reported results related to acquisition are shown only for experiments in which exercise was performed before motor task practice. Reported retention test results are shown regardless of whether exercise was performed before or after motor task practice. (Panel a) Reported results from acquisition. (Panel b) Reported results on motor learning from retention tests conducted within 24 hours following exercise, (Panel c) Reported results on motor learning from retention tests conducted at 24 hours following exercise (Panel d) Reported results on motor learning from retention tests conducted after 24 hours following exercise. Green dots indicate low-intensity exercise, orange dots indicate moderate-intensity exercise, and red dots indicate high-intensity exercise.

#### 3.5.1 Skill acquisition

A total of 49 experimental conditions (E = 49) assessed the effects of physical exercise on motor skill acquisition (see **Figure 6a**). In healthy children (e = 3; all high intensity), no significant effects were observed. In healthy adults (e = 35), the majority of experimental conditions showed no effect (e = 28; across low, moderate, and high intensities), while six conditions reported enhancement (e = 6; three with high intensity, three with moderate intensity), and one moderate-intensity condition led to attenuated performance. In healthy older adults (e = 3), two conditions showed no effect (high and moderate intensity), and one high-intensity condition led to attenuated performance. Among individuals with stroke (e = 4), one moderate-intensity condition enhanced acquisition, while the others (two high intensity and one moderate intensity) showed no effect. In individuals with Parkinson’s disease, three conditions (e = 3; 2 moderate and 1 high intensity) reported no difference. Finally, the single experimental condition involving individuals with Huntington’s disease (e = 1) reported improvement following a moderate-intensity exercise.

#### 3.5.2 Retention tests less than 24h

A total of 28 experimental conditions (E = 28) assessed the effects of physical exercise on performance within 24 hours following the exercise intervention (see **Figure 6b**). In healthy children (e = 3), two experimental conditions showed enhanced performance (both at high intensity), and one condition at high intensity showed no difference. In healthy adults (e = 23), five conditions demonstrated enhanced performance (three at high intensity, two at moderate intensity), 17 showed no difference (across all three intensities), and one high-intensity condition showed attenuated performance at retention. In healthy older adults (e = 2), one condition showed enhanced performance (high intensity) and one showed no difference (moderate intensity). Notably, no experimental conditions involving clinical populations were reported in this retention time category.

#### 3.5.3 Retention tests at 24h

A total of 49 experimental conditions (E = 49) assessed the effects of physical exercise on performance at 24 hours following the exercise intervention (see Figure 6c). In healthy children (e = 4), three conditions showed enhanced performance (all at high intensity), and one condition showed attenuated performance (high intensity). In healthy adults (e = 36), 12 conditions demonstrated enhanced performance (nine at high intensity, two at moderate intensity, and one at low intensity), while 24 conditions showed no difference (across all intensities). In healthy older adults (e = 2), both conditions showed no difference (one at high intensity and one at moderate intensity). Among individuals with stroke (e = 4), one condition showed enhanced performance (high intensity), and three conditions showed no difference (all at high intensity). In individuals with Parkinson’s disease (e = 3), one condition showed enhanced performance (moderate intensity), and two conditions showed no difference (one at high intensity and one at moderate intensity).

#### 3.5.4 Retention tests more than 24h

A total of 31 experimental conditions (E = 31) assessed the effects of physical exercise on performance beyond 24 hours following the exercise intervention (see **Figure 6d**). In healthy children (e = 3), all conditions showed enhanced performance (all at high intensity). In healthy adults (e = 21), fourteen conditions showed enhanced performance (ten at high intensity, two at moderate intensity, and two at low intensity), and seven conditions showed no difference (three at high intensity, three at moderate intensity, and one at low intensity). In healthy older adults one condition showed no difference (high intensity). Among individuals with stroke (e = 2), one condition showed no difference (moderate intensity), and one condition showed attenuated performance (moderate intensity). In individuals with Parkinson’s disease (e = 3), all conditions showed enhanced performance (two at high intensity and one at moderate intensity).

### 3.6 Review articles previously published

Among the included sources, 4 secondary sources were review articles. Bonuzzi and colleagues conducted a systematic review on aerobic exercise and motor learning in healthy individuals, highlighting that exercise performed before or after motor task practice can improve motor learning (Bonuzzi & Torriani-Pasin, 2022). Hubner and colleagues conducted a systematic review examining the effects of physical activity level and acute bouts of physical exercise on motor performance and motor learning of upper limb tasks in older people. This systematic review highlighted the benefits of a high physical activity level in the early phases of motor learning (Hubner & Voelcker-Rehage, 2017). Taubert and colleagues performed a narrative review and discussed behavioral and neurobiological evidence in favor of aerobic exercise as a neuromodulator strategy to promote motor learning and neuroplasticity (Taubert et al., 2015). Wanner and colleagues performed a systematic review and meta-analysis and evaluated how acute aerobic exercise affects the encoding and consolidation of motor memory, highlighting the importance of timing and intensity (Wanner et al., 2020b).

## 4. Discussion

To the best of our knowledge, this is the first scoping review to explore and map the existing literature on the use of physical exercise to prime motor learning. We included studies involving any population type, any form of physical exercise, and whether the physical exercise was performed before or after any motor learning task. Although our scoping review focuses largely on experimental studies (62 included studies), it should be noted that four previous reviews have explored specific aspects of the topic (1 narrative review, 2 systematic reviews and 1 systematic review with meta-analysis). These focused reviews each addressed important aspects of the topic, and our scoping review synthesis builds on and integrates these contributions by examining a broader range of populations, exercise types, motor learning categories and exercise timing in a single comprehensive framework.

The main outcomes of this scoping review were the following: (1) the evaluated population consisted primarily of healthy individuals, with limited focus on clinical populations and children; (2) the predominant type of physical exercise used was lower-limb cycling, potentially due to its ease of implementation; (3) resistance exercise was rarely used, highlighting a significant gap in the literature regarding its potential role in the promotion of motor learning; (4) in several studies, the prescribed exercise intensity did not align accurately with ACSM guidelines, which may limit the ability to draw accurate conclusions about the effects of specific exercise intensities on motor learning; (5) many studies conducted a single retention test on the same day as the practice session; however, according to the classical definition of motor learning, demonstrating maintenance in motor skill improvement over time via delayed retention tests is essential (Schmidt et al., 2018), and hence these findings may not truly reflect motor learning.

### 4.1 Limited representation of clinical populations

Most of the included studies involved healthy individuals, with children and clinical populations being significantly less evaluated. One reason could be due to practical limitations, as recruiting healthy adult participants is usually less complicated and requires less time and resources. Yet, in order to determine whether the results obtained in healthy individuals are transferable to children and individuals living with specific health-related conditions, it is essential to replicate the interventions in children and clinical populations. This is particularly of interest for children as this population is crucially developing motor skills needed for their adult life. It is also crucial for clinical populations given that many interventions improving motor learning are designed for rehabilitation purposes (Krakauer, 2006), and the clinical relevance of these interventions should be tested within the clinical populations of interest.

Furthermore, given the limited number of studies involving clinical populations, the division across three distinct conditions (stroke, Parkinson’s disease, and Huntington’s disease) further limits the applicability of the findings to specific clinical settings, as each subgroup is represented by only a small number of studies.

Importantly, the underlying mechanisms of motor learning may differ between healthy individuals and clinical populations due to different factors including impaired neuroplasticity, altered sensory feedback, or compensatory motor strategies (Boyd & Winstein, 2006; Buma et al., 2010). For instance, in individuals with stroke, cortical reorganization and the recruitment of alternative neural pathways may affect how motor skills are re-acquired (Nudo, 2006). Similarly, in Parkinson’s disease, basal ganglia and sensorimotor-related dysfunction may alter feedback-based learning mechanisms (Boyd et al., 2004). These differences imply that exercise interventions that are effective in healthy individuals may not produce the same effects in the context of clinical conditions, making it essential to investigate tailored strategies specific to each clinical population. Therefore, prioritizing research in populations with motor dysfunction and rehabilitation needs including individuals with stroke, or other populations living with neurodegenerative disorders, would be particularly valuable. These population groups could benefit the most from interventions that can enhance motor learning, recovery and maintenance of functional independence. By targeting clinical populations, we can develop more effective, evidence-based interventions that address their specific motor learning profiles.

### 4.2 Types of aerobic exercise: a focus on lower-limb cycling and the need for diversity

Among the included studies, the predominant form of aerobic exercise was lower-limb cycling. This type of aerobic exercise is likely favored due to its ease of implementation, low risk of injury, and practicality in experimental settings. Particularly, lower-limb cycling effectively induces an aerobic response by engaging the leg muscles, the largest muscle group in the body, resulting in a substantial cardiovascular activation (Poole & Richardson, 1997; Scott et al., 2021). Also, lower-limb cycling allows the upper limbs to remain at rest, which helps prevent the development of neuromuscular fatigue that could negatively impact motor learning when upper-limb motor tasks are used (Ranieri & Di Lazzaro, 2012).

Other types of aerobic exercise such as running, walking, dancing and upper-limb cycling were considerably less represented. Including these diverse forms of aerobic exercise in future research is important for several reasons. Different types may activate distinct neuromuscular and cardiovascular responses, influencing task-specific adaptations, and offer practical alternatives for various populations.

For example, running engages more complex coordination, while upper-limb cycling may be particularly suitable for individuals with lower-limb limitations. Dancing, which is often practiced in groups and especially popular among older adults also deserves attention, as it not only combines demands on the aerobic system and coordination (Keogh et al., 2009; Limanskaya et al., 2021), but has also been associated with enhanced cognitive function (Hewston et al., 2021; Karpati et al., 2015). Dancing also may boost motivation and adherence to regular engagement in physical activity through social interaction (Brustio et al., 2018). In addition, music, which is an integral part of most dance interventions, has itself been linked to improved cognitive function (Schellenberg, 2005) and positive affect (Campion & Levita, 2014), further supporting the potential of dance-based activities in promoting motor learning. Such variety of physical exercise could help reveal how the specific demands of different types of exercise might affect motor learning.

Importantly, all of these reported aerobic types of physical exercise primarily relied on concentric muscle contractions, where the muscles shorten while producing a force. None of the included studies investigated aerobic exercise involving primarily eccentric contractions, like eccentric cycling, in which muscles lengthen while producing a force.

### 4.3 Eccentric and resistance exercise: underexplored avenues in motor learning research

Interestingly, none of the included studies investigated the effects of physical exercise primarily involving eccentric muscle contractions on motor learning. Eccentric contractions, which involve muscle lengthening while resisting a workload, are known to induce distinct neural adaptations compared to concentric contractions (Fang et al., 2001; Latella et al., 2019). Despite this, research focusing specifically on eccentric-based physical exercise remains limited, representing a notable gap in the literature. Specifically, eccentric contractions have been associated with a prolonged decrease in intracortical inhibition (Latella et al., 2019), a neurophysiological marker of altered motor cortex excitability. Also, it has been shown that eccentric cycling elicits greater activation in sensorimotor-related cortical areas of the frontal and parietal lobes during movement planning and execution (Fang et al., 2001), as well as increased activation in cognitive-related brain regions (Borot et al., 2024). Given the involvement of these brain regions in motor learning, eccentric cycling could induce potential benefits to motor learning.

In fact, recent preliminary evidence from our laboratory (Youssef et al., 2025) supports this idea. This study, which was not included in the present scoping review due to its preprint status at the time of submission, suggests that eccentric cycling enhances motor learning more than concentric cycling in healthy young adults. These findings suggest that compared to concentric cycling, eccentric cycling may offer specific advantages that promote motor learning. Beyond its potential effects on motor learning, eccentric cycling also provides physiological advantages. For a similar workload, eccentric cycling results in a lower cardiovascular, respiratory, and metabolic cost compared to concentric cycling (Clos et al., 2019; Isner-Horobeti et al., 2013; Peñailillo et al., 2013; Peñailillo et al., 2015). Eccentric cycling is therefore a particularly appealing exercise modality for clinical populations with reduced cardiorespiratory capacity [e.g., (Pageaux et al., 2020)], as well as for older adults [(e.g., (Gault & Willems, 2013)]. Since lower-limb cycling is already the most employed exercise type in the reviewed literature, integrating eccentric cycling could allow researchers to build on already established protocols while expanding the range of muscular contractions studied. Finally, other eccentric-focused exercise types like downhill walking or running also deserve attention. These exercise types involve distinct neuromuscular demands (Garnier et al., 2018) and a lower energy cost (Vernillo et al., 2017), which could be beneficial in rehabilitation. Given the potential benefits of eccentric exercise on motor learning, its integration into future studies evaluating the effects of physical exercise on motor learning appears both feasible and promising (LaStayo et al., 2014).

Beyond eccentric-focused types of exercise, our findings also raise important considerations about resistance exercise, which often incorporates eccentric contractions and shares similar physiological mechanisms (Douglas et al., 2017). These findings also highlight a significant gap in the motor learning literature, which is the underrepresentation of resistance exercise as priming exercise. Of the 81 experimental conditions included, only two involved resistance exercise, despite the fact that this exercise type is widely used in rehabilitation and training. Resistance exercise may influence motor learning through several neural mechanisms. It has been shown that resistance exercise increases muscle activation and neural drive to muscles, leading to improved force production capacity (Aagaard et al., 2002). Resistance exercise can also modulate corticospinal excitability, with evidence indicating that resistance training induces neuroplastic changes within the motor cortex and corticospinal tract (Carroll et al., 2002; Kidgell & Pearce, 2010). Furthermore, resistance exercise may enhance proprioceptive input, increasing sensory feedback, which is an essential component of motor control and learning (Proske & Gandevia, 2012). Supporting these observations, cerebral blood flow during force exertion has been shown to correlate with activity in brain areas such as the primary motor cortex, and the supplementary motor area (Dettmers et al., 1995). These regions are also integral to motor learning processes, suggesting a neuroanatomical overlap between areas recruited during resistance exercise and those underlying motor learning. However, the direct role of the effects of resistance exercise on motor learning remains largely unexplored. Investigating how resistance exercise influences motor learning represents a promising avenue for future research. Taken together, these observations highlight the need for future studies to incorporate both eccentric and resistance exercise types to better understand their potential roles in enhancing motor learning across various populations.

### 4.4 Prescription of exercise intensities

Although all studies reported the exercise intensity used, we standardized intensity categorization based on the exercise prescription details provided using the ACSM guidelines (ACSM, 2018). In 5 experimental conditions, discrepancies were observed between the prescribed intensity and the actual workload performed by participants. Some exercise appeared either lighter or more intense than prescribed. For instance, Holman et al. (2021) and Statton et al. (2015) reported using moderate-intensity exercise, yet the prescribed workloads correspond to high-intensity levels according to ACSM guidelines (Holman & Staines, 2021; Statton et al., 2015). Similarly, Opie and Semmler (2019) described their exercise protocol as low-intensity, although it aligns with moderate-intensity based on ACSM classifications (Opie & Semmler, 2019). In contrast, Loras et al. (2020) referred to their two exercise conditions as moderate and high intensity exercise, but these would correspond to low and moderate intensity, respectively, under ACSM classifications (Loras et al., 2020). First, this inconsistency suggests that following established exercise guidelines is essential for drawing accurate conclusions about the effects of specific exercise intensities on motor learning. Second, it allows replicability across studies which is an essential factor for building a consistent and reliable body of evidence. Additionally, while many studies relied on predicted values (e.g., age-predicted maximal heart rate, age-predicted heart rate reserve, estimated VO2max), and although we recognize the practicality of using predicted values in certain research settings, prescribing intensities based on measured capacities are preferable to ensure that the targeted intensity is truly achieved for each participant. Interestingly, none of the studies included in this review employed self-paced exercise protocols. This is notable, as self-paced exercise, for example prescribed with the intensity of perceptual responses such as effort (Beaume et al., 2025; Pageaux, 2016), when matched for workload with externally prescribed intensities (Ekkekakis et al., 2008), has been proposed to enhance affective responses, which can be beneficial for promoting adherence to rehabilitation programs (Ekkekakis et al., 2011).

### 4.5 Uneven representation of motor learning categories

While the studies included in this review examined various motor learning categories such as sequence learning, motor acuity, and motor adaptation, none investigated the effects of physical exercise on associative learning, which is a fundamental form of motor learning involving stimulus-response associations. Given its relevance in real-world contexts and rehabilitation settings (Le Pelley et al., 2016), investigating how physical exercise influences associative learning could offer valuable insights. For instance, tasks such as learning to respond appropriately to traffic signals while driving rely heavily on stimulus-response associations.

Additionally, the field would benefit from more balanced research across motor learning categories. The current emphasis on motor acuity (n = 30) contrasts with the limited number of studies evaluating sequence learning and motor adaptation (both n = 16). Sequence learning, for example, is highly relevant for activities like playing a musical instrument or typing on a keyboard, while motor adaptation is crucial in situations such as adjusting movements to a new sports technique or learning to walk on uneven surfaces. More research in these underrepresented categories could allow for more robust meta-analyses and potentially clarify whether the effects of physical exercise vary depending on the motor learning type.

### 4.6 Timing of retention tests

When measuring skill retention, different time points were reported across the studies falling in our defined time windows: less than 24 hours, at 24 hours, and more than 24 hours following motor task practice. Some studies included two time points (Chan et al., 2023; Charalambous et al., 2019; Dal Maso et al., 2018; Hubner et al., 2018; Jespersen et al., 2023; Khan et al., 2022; Pixa et al., 2021; Thomas et al., 2017; Thomas et al., 2016; Wanga et al., 2023; Wanner et al., 2021; Yildirim et al., 2024) or all three time points (Angulo-Barroso et al., 2019; Ferrer-Uris et al., 2018; Ferrer-Uris et al., 2017; Holman & Staines, 2021; Lundbye-Jensen et al., 2017; Roig et al., 2012; Skriver et al., 2014; Swarbrick et al., 2020; Tomporowski & Pendleton, 2018), which provides a clearer understanding of the effects on motor learning. Other studies only assessed short-term retention at less than 24 hours (Bartz & Smith, 1970; Bosch et al., 2020; Coco et al., 2016; James & Wang, 2023; Kuo et al., 2023a; Kuo et al., 2023b; Ostadan et al., 2016; Stavrinos & Coxon, 2017; Taylor et al., 2024). However, measuring skill retention only in the short term (e.g., same day as initial practice) may not be sufficient to assess motor learning, as this may primarily reflect temporary rather sustained performance improvements (Schmidt et al., 2018). According to Schmidt et al. (2018), observing performance after a sufficient delay provides stronger evidence of stable and relatively permanent changes in the learner’s capability, which is the hallmark of motor learning (Schmidt et al., 2018).

Ideally, studies could include both short-term and longer-term retention tests, with the 24-hour time point used as a standard. This would help determine whether performance improvements due to practice are maintained over time, facilitate comparisons across studies, and provide insight into the evolution of motor skill consolidation, which is the process by which newly acquired motor skills become stable and resistant to interference. Using consistent methods to measure skill retention would also help improve the quality of future studies in the field of physical exercise and motor learning.

### 4.7 Limitations

This scoping review has several limitations, particularly regarding the reported results of motor learning, due to the complexity of categorization. First, for retention tests, all studies were grouped together regardless of whether physical exercise was performed before or after skill acquisition, as the heterogeneity in study designs made it difficult to establish clear subcategories for the reported results on motor learning. Second, no differentiation was made based on the type of exercise for the reported results on motor learning, which limits the ability to attribute specific effects to a particular type of exercise. Third, because we examine the effects of physical exercise in general and as reported by the authors, exercise frequency was not considered, and no distinction was made between acute and chronic interventions. Future studies with more data may allow for a more refined mapping of the literature addressing these aspects.

### 4.8 Perspectives for future studies

This scoping review highlighted several key areas for future research on the effects of physical exercise on motor learning. First, accurately prescribing exercise intensity is essential. Many studies showed inconsistencies between the reported and the actual prescribed intensity, which can affect the reliability of findings. Using measured values rather than predicted ones, and adhering to established guidelines like the ACSM, would improve consistency and accuracy across studies. As self-paced exercise are proposed to enhance affective responses (Beaume et al., 2025; Pageaux, 2016) and have positive effect on exercise adherence, future studies should consider testing the possibility to prime motor learning with exercise prescribed based on perceptual responses, with particular interest on the perception of effort (Beaume et al., 2025; Pageaux, 2016). Furthermore, although most studies have concentrated on concentric lower-limb cycling, it is essential to explore the impact of other types of exercise, such as eccentric cycling, on motor learning. Additionally, resistance exercise was rarely used. Integrating resistance exercise modalities will enable us to better understand its potential role in enhancing motor learning across various populations. Moreover, measuring skill retention in motor learning studies in exercise science is another area to improve. Studies testing retention only within a few hours may not fully capture the achievement and maintenance of motor learning. Therefore, including longer retention time points, particularly at 24 hours and more, might help in providing a more comprehensive understanding of the effects of physical exercise on motor learning. Finally, more studies are needed where children, older adults and clinical populations are involved. Increasing research on clinical populations is especially important as motor learning can play a critical role in rehabilitation and recovery.

## 5. Conclusion

In conclusion, while the field of physical exercise and motor learning is growing, there are several key areas that require improvement. Future research should prioritize accurate and consistent reporting of exercise intensity and consider longer retention intervals to better capture lasting motor learning effects and, consequently, better understand the impact of physical exercise on motor learning. Additionally, greater inclusion of children, older adults and clinical populations is essential, as these population groups may benefit most from motor learning interventions but remain underrepresented in the current literature. Expanding the range of exercise modalities, more particularly types of exercise involving eccentric contractions and resistance-based exercise, may also reveal unique advantages relevant to motor learning in both healthy and clinical populations. Addressing these gaps will help in building a more robust and applicable body of evidence to inform both clinical and non-clinical practice.

## List of Abbreviations

n =: subtotal number of participants
k =: subtotal number of studies
g =: subtotal number of groups
e =: subtotal number of experimental conditions
N =: total number of participants (grand total)
K =: total number of studies (grand total)
G =: total number of groups (grand total)
E =: total number of experimental conditions (grand total)§

## Author contributions

**Layale Youssef:** Conceptualization, Methodology, Formal analysis, Investigation, Writing – Original Draft, Review & Editing, Visualization. **Amanda O’Farrell:** Formal analysis, Investigation, Writing – Review & Editing**. Denis Arvisais:** Methodology, Investigation, Writing – Review & Editing**. Jason L. Neva:** Conceptualization, Writing – Review & Editing, Visualization, Supervision**. Benjamin Pageaux:** Conceptualization, Writing – Review & Editing, Visualization, Supervision.

## Supporting information

Data_Extraction_Sheet

## Supplementary Material 1

**Figure S1.**
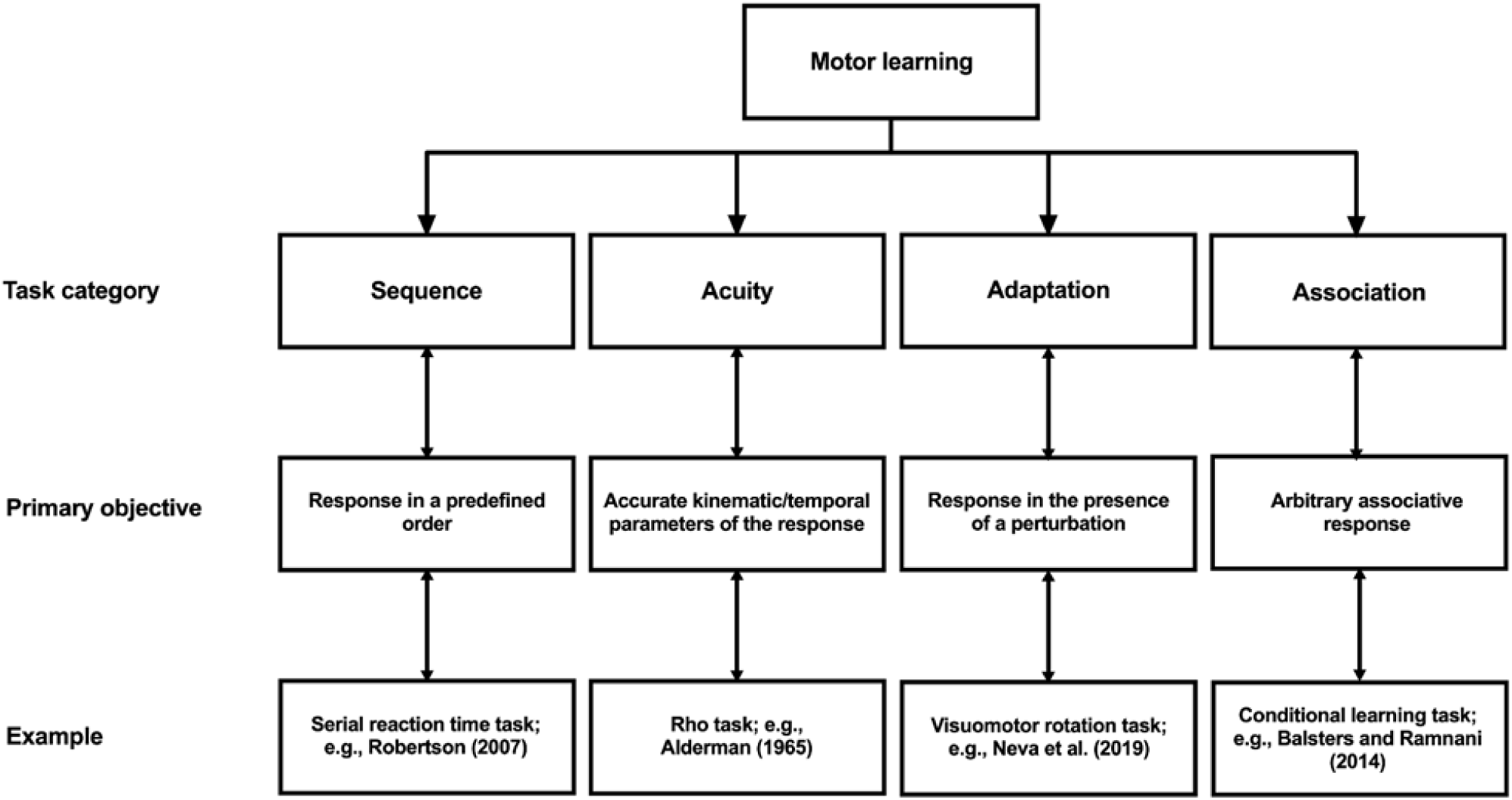
Classification of the task categories investigated to infer motor learning. This figure illustrates four commonly used motor learning task categories, classified according to their primary objective. The examples shown represent typical experimental tasks used within each category to assess motor learning.

---

**Table S1:** Characteristics of the studies included in the scoping review.

| Study | Total N | Type of population (n of exercise groups; age ± SD years) | Physical exercise | Exercise parameters | Motor Learning Category | Exercise Timing to Acquisition | Retention | Exercise effect on motor learning compared to control | |
| --- | --- | --- | --- | --- | --- | --- | --- | --- | --- |
|  |  |  |  |  |  |  |  | Acquisition | Retention |
| Andrews et al., 2023 | 20 | Huntington's disease (n = 10, 48.4 ± 13) | Lower-limb cycling | Moderate<br>20 min - CONT<br>50-55% age-predicted HRR | Motor acuity | Before | 7d | ↗ | ↔ |
| Andrushko et al., 2023 | 25 | Stroke patients (n = 14, 66 ± 11) | Lower-limb cycling (5 sessions) | High<br>23 min - HIIT<br>3min at 75% Wpeak<br>3 min at 10W | Sequence | Before | 24h | ↔ | ↔ |
| Angulo-Barroso et al., 2019 | 71 | Healthy children (n = 46, 9.2 ± 0.9) | Running | High<br>16 min (LONG) : 5 min (SHORT) - HIIT<br>LONG: 3 min at 85% estim. VO<sub>2</sub>max ;<br>2 min at 60% estim. VO<sub>2</sub>max ;<br>SHORT: 2 min at 85% estim. VO<sub>2</sub>max ;<br>1 min at 60% estim. VO<sub>2</sub>max | Motor adaptation | Before | 1h, 24h, 7d | ↔ | ↗ for LONG and SHORT |
| Baird et al., 2018 | 48 | Healthy adults (n = 32, 23 ± 3.1) | Lower-limb cycling | Low<br>CONT: 40% measured max resistance until 200 kals were expended (average: 28.67 min)<br>High<br>CONT: 80% measured max resistance until 200kals were expended (average 16.75 min) | Sequence | Before | 24h | ↔ | ↔ |
| Bartz et al., 1970 | 20 | Healthy adults (n = 10, 18-20 yrs) | Walking | Moderate<br>4 min - CONT<br>HR target: 120-170 bpm | Motor adaptation | Before | <10 min | ↔ | ↔ |
| Bonuzzi et al., 2023 | 90 | Stroke patients (n = 15; 51.6 ± 9.2) | Lower-limb cycling (3 sessions) | Moderate<br>20 min - CONT<br>50-70% age-predicted HRR | Sequence | Before | 10d | ↔ | ↘ |
| | | Stroke patients (n = 15; 49.6 ± 15.9) | | | | After | | ↔ | ↔ |
| | | Healthy adults (n = 15; 51.6 ± 9.2) | | | | Before | | ↘ | ↔ |
| | | Healthy adults (n = 15; 49.6 ± 15.9) | | | | After | | ↔ | ↔ |
| Bosch et al., 2020 | 15 | Healthy adults (n = 15; 23.7 ± 4.0) | Lower-limb cycling | Moderate<br>30 min - CONT<br>70% measured MHR<br>High<br>15 min - CONT<br>80% measured MHR | Sequence | After | 2.5h | NA | ↗ for High<br>↔ for Moderate |
| Chan et al., 2023 | 30 | Parkinson's disease (n = 15; 62.4 ± 8.5) | Lower-limb cycling | High<br>20 min - CONT<br>60-75% measured HRR | Sequence | After | 1d, 7d | ↔ | ↗ at Day 7 |
| Charalambous et al., 2018 (active control) | 37 | Stroke patients (n=12; 55 ± 16) | Treadmill walking | High<br>5 min - CONT<br>70-85% age-predicted MHR | Motor adaptation | Before | 24h | ↔ | ↔ |
| | | Stroke patients (n=12; 62 ± 10) | Total body exercise | | | After | | ↔ | ↔ |
| Charalambous et al., 2019 | 26 | Healthy adults (n = 13; 21.8 ± 0.5) | Whole-body cycling | High<br>5 min - CONT<br>77-94% age-predicted MHR | Motor adaptation | Before | 1d, 7d | ↔ | ↔ |
| Chen et al., 2023 | 24 | Healthy adults (n = 12; 20.9 ± 0.7) | Treadmill Running | Moderate<br>30 min - CONT<br>65-85% age-predicted MHR (average was 75%) | Motor acuity | Before | 24h | ↔ | ↔ |
| Coco et al., 2016 (no control) | 28 | Healthy adults (n = 28; 37.4 ± 5.8) | Lower-limb cycling | High<br>CONT: 3 minutes of unloaded cycling with 30W increase every 3min until exhaustion | Motor acuity | After | < 1h | NA | ↘ |
| Cristini et al., 2023 | 56 | Healthy adults (n = 20; 23.9 ± 3.4) | Lower-limb cycling | High<br>16 min - HIIT<br>3 min at 85-90 % of measured PPO<br>2min at 25 % of PPO. | Sequence | After | 24h | NA | ↔ |
| Dal Maso et al., 2018 | 25 | Healthy adults (n = 12; 24.3 ± 3.6) | Lower-limb cycling | High<br>15 min - HIIT<br>3min at 90% measured HRR<br>2min at 50W | Motor acuity | After | 8h, 24h | NA | ↔ for 8h<br>↗ at 24h |
| Duchesne et al., 2015 | 39 | Parkinson's disease (n = 19; 59 ± 7.1) | Lower-limb cycling (3x/week for 12 weeks) | High<br>CONT<br>Start: 20 min at 60% measured PPO<br>End: 40 min at 80% measured PPO | Sequence | After | 12 weeks | NA | ↗ |
| | | Healthy older adults (n = 20; 64 ± 8.2) | | | | After | | | ↗ |

CONT = continuous; HRR = Heart rate reserve; W = Watts; HIIT = high-intensity interval training; kals = kilocalories; bpm = beats per minute; HR = Heart Rate; MHR = maximal heart rate; PPO = peak power output; estim. = estimated; Before = exercise was performed before the motor task practice; After = exercise was performed after the motor task practice; min = minutes; h = hours; d = days; NA = not available; ↗ = enhancement following exercise as reported by the authors; ↘ = decrease following exercise as reported by the authors; ↔ = no change following exercise as reported by the authors.

**Table S1. continued**
| Study | Total N | Type of population (n of exercise groups; age ± SD years) | Physical exercise | Exercise parameters | Motor Learning Category | Exercise Timing to Acquisition | Retention | Exercise effect on motor learning compared to control | |
| --- | --- | --- | --- | --- | --- | --- | --- | --- | --- |
|  |  |  |  |  |  |  |  | Acquisition | Retention |
| Ferrer-Uri et al., 2017 | 29 | Healthy adults (n = 10, 20.9 ± 1.8) | Shuttle run | High<br>13 min - HIIT<br>3min at 85% estimated VO<sub>2</sub>max<br>2min at 60% estimated VO<sub>2</sub>max | Motor adaptation | Before | 1h, 24h, 7d | ↔ | ↗ at 1h |
| | | Healthy adults (n = 10, 20.5 ± 1.8) | | | | After | | NA | ↗ at 1h |
| Ferrer-Uri et al., 2018 | 33 | Healthy children (n = 10, 9.2 ± 1.1) | Shuttle run | High<br>13 min - HIIT<br>3min at 85% estimated VO<sub>2</sub>max<br>2min at 60% estimated VO<sub>2</sub>max | Motor adaptation | Before | 1h, 24h, 7d | ↔ | ↗ |
| | | Healthy children (n = 12, 9.1 ± 0.8) | | | | After | | NA | ↔ |
| Greeley et al., 2021 | 44 | Healthy older adults (n = 19; 68.3 ± 9.2) | Lower-limb cycling (5 sessions) | High<br>23 min - HIIT<br>3min at 75% measured Wpeak<br>3min at 10W | Sequence | Before | 24h, 35 days | ↔ | ↔ |
| Helm et al., 2017 | 54 | Healthy adults (n = 27; 24.5 ± 2.8) | Upper-limb cycling | High<br>5 min - HIIT<br>1 min at 80% age-predicted MHR<br>1 min at half of cycling power | Motor adaptation | Before | 24h | ↔ | ↔ |
| Holman et al., 2021 | 33 | Healthy adults (n = 17; 21.6 ± 2.3) | Lower-limb cycling | High<br>5 min - CONT<br>70% age-predicted HRR | Motor adaptation | After | 30min, 24h, 7d | NA | ↗ at 7d |
| Hubner et al., 2018 | 38 | Healthy older adults (n = 17; 68.2 ± 3.2) | Lower-limb cycling | Moderate<br>25 min - CONT<br>60% measured Wpeak | Motor acuity | Before | 30 min, 24h | ↔ | ↔ |
| Hung et al., 2021 | 44 | Healthy adults (n = 22; 23 ± 2) | Lower-limb cycling | High<br>25 min - HIIT<br>1 min at 90% measured Wpeak ;<br>1 min 25% measured Wpeak | Motor acuity | After | 24h | NA | ↔ |
| James and Wang., 2023 | 16 | Healthy adults (n = 16; 24.6 ± 4.7) | Lower-limb cycling | Moderate<br>25 min - CONT<br>65-70% age-predicted MHR | Motor adaptation | Before | < 30min | ↔ | ↔ |
| Jespersen et al., 2023 | 64 | Healthy adults (n = 16; 24.1 ± 3.9) | Lower-limb cycling | Moderate<br>20 min - HIIT<br>3min at 45% measured Wpeak ;<br>2 min at 50W | Sequence | Before | <5 min, 7d | ↔ | ↗ at 7d |
| | | Healthy adults (n = 16; 23.8 ± 4.2) | | High<br>20 min - HIIT<br>3min at 90% measured Wpeak ;<br>2 min at 50W | | After | | NA | ↗ at 7d |
| Khan et al., 2022 | 48 | Healthy adults (n = 23; 22.9 ± 4.9) | Lower-limb cycling | High<br>20 min - HIIT<br>3min at 90% measured Wpeak<br>2min at 50W | Motor acuity | After | 24h, 7d | NA | ↗ at 7d |
| Kuo et al., 2023a | 26 | Healthy adults (n = 26; 27.2 ± 1.0) | Lower-limb cycling | Low<br>20 min - CONT<br>40-60% age-predicted MHR<br>Moderate<br>20 min - CONT<br>61-74% age-predicted MHR<br>High<br>20 min - CONT<br>75-95% age-predicted MHR | Sequence | After | 0-5 min | NA | ↗ for Moderate |
| Kuo et al., 2023b | 20 | Healthy adults (n = 20; 26.4 ± 1.0) | Lower-limb cycling | Moderate<br>20 min - CONT<br>61-74% age-predicted MHR | Sequence | After | 0-5 min | NA | ↗ |
| Lehmann et al., 2020 | 31 | Healthy adults (n = 15; 23 ± 7) | Lower-limb cycling (7 sessions) | High<br>20 min - HIIT<br>2x3min at 170 (PWC)<br>4 min at 120 (PWC) | Motor adaptation | Before | 8 weeks | ↗ | ↗ |
| Loras et al., 2020 (no control) | 40 | Healthy adults (n = 40; 23.8 ± 2.0) | Lower-limb cycling | Low<br>25 min - CONT<br>50% age-predicted MHR<br>Moderate<br>25 min - CONT<br>75% age-predicted MHR | Motor acuity | Before | 24h | ↔ | ↔ |
| Lundbye-Jensen et al., 2017 | 77 | Healthy children (n = 26; 10.4 ± 0.7) | Floorball | High<br>15 min - HIIT<br>3 min at high intensity<br>2 min at low intensity<br>(measured via HR monitoring) | Motor acuity | After | 1h, 24h, 7d | ↔ | ↗ at 7d |
| | | Healthy children (n = 25; 10.8 ± 0.8) | Running | | | After | | | ↘ at 1h<br>↗ at 7d |
| Mang et al., 2014 | 16 | Healthy adults (n = 16; 23.9 ± 3.7) | Lower-limb cycling | High<br>20 min - HIIT<br>3min at 90% measured PPO<br>2min at 50W | Sequence | Before | 24h | ↗ | ↗ |
| Mang et al., 2016 | 16 | Healthy adults (n = 16; 25.7 ± 4.6) | Lower-limb cycling | High<br>20min - HIIT<br>3min at 90% measured PPO<br>2min at 50W | Sequence | Before | 24h | ↔ | ↗ |

**Table S1. continued**
| Study | Total N | Type of population (n of exercise groups; age ± SD years) | Physical exercise | Exercise parameters | Motor Learning Category | Exercise Timing to Acquisition | Retention | Exercise effect on motor learning compared to control | |
| --- | --- | --- | --- | --- | --- | --- | --- | --- | --- |
|  |  |  |  |  |  |  |  | Acquisition | Retention |
| Munz et al., 2021 | 12 | Healthy children (n = 12; 10.2 ± 1.4) | Lower-limb cycling | High<br>40 min - CONT<br>(4 blocks of 10 min)<br>85-90% of age-predicted MHR | Motor acuity | After | 24h | NA | ↗ |
| Nepveu et al., 2017 | 22 | Stroke patients (n = 11; 64.7 ± 11.6) | Lower-limb cycling | High<br>15 min - HIIT<br>3 min at 100% measured W<sub>peak</sub><br>2min at 25% measured W<sub>peak</sub> | Motor acuity | After | 24h | NA | ↗ |
| Neva et al., 2019 | 17 | Healthy adults (n = 17; 24 ± 3) | Lower-limb cycling | Moderate<br>25 min - CONT<br>65-70% age-predicted MHR | Motor adaptation | Before | 24h | ↗ | ↗ |
| Ostadan et al., 2016 | 38 | Healthy adults (n = 25 ; 22.4 ± 3.9) | Lower-limb cycling | High<br>15 min - HIIT<br>3 min at 85-90% measured VO<sub>2</sub>peak<br>2 min at 25% measured W<sub>peak</sub> | Sequence | After | 8h | NA | ↔ |
| Opie and Semmler., 2019 | 13 | Healthy adults (n = 13 ; 24.0 ± 3.3) | Lower-limb cycling | Moderate<br>30 min - CONT<br>50% age-predicted HRR<br>High<br>30 min - HIIT<br>77% age-predicted HRR | Motor acuity | Before | 24h | ↗ for Mod | ↗ for Mod |
| Perini et al., 2016 | 38 | Healthy adults (n = 18; 22.9 ± 2.2) | Lower-limb cycling | Moderate<br>30 min - CONT<br>70% measured MHR | Motor acuity | Before | NA | ↗ | NA |
| Pixa et al., 2021 | 27 | Healthy adults (n = 30; 23.7 ± 4.4) | Lower-limb cycling | Low<br>25 min - CONT<br>20% measured PPO<br>High<br>23 min - HIIT<br>3 min at 90% measured W<sub>peak</sub><br>2 min at 60% measured W<sub>peak</sub> | Motor acuity | Before | 15 min, 30min, 24h | ↔ | ↔ |
| Quaney et al., 2009 | 21 | Stroke patients (n = 19; 64.1 ± 12.3) | Lower-limb cycling (24 sessions) | Moderate<br>45 min - CONT<br>70% measured MHR | Sequence | Before | 8 weeks | ↗ | ↔ |
| Quinlan et al., 2021 | 29 | Healthy adults (n = 15; 45 ± 5) | Lower-limb cycling | High<br>20 min - HIIT<br>4 min at 90% measured W<sub>peak</sub><br>2min of active recovery at 50-80W | Motor acuity | Before | 5-7d | ↔ | ↔ |
| Rhee et al., 2016 | 60 | Healthy adults (n = 22; 19.8 ± 1.3) | Lower-limb cycling | High<br>21 min - CONT<br>18min at 80% measured MHR<br>3min at 100% measured MHR | Motor acuity | After | 24h | NA | ↗ |
| Roig et al., 2012 | 48 | Healthy adults (n = 16; 24.1 [21-33]) | Lower-limb cycling | High<br>15 min - HIIT<br>3min at high intensity<br>2min at 50W<br>Based on measured lactate level<br>Workload (200-315) | Motor acuity | Before | 1h, 24h, 7d | ↔ | ↗ for 24h and 7d (higher when exercise is performed after) |
|  |  | Healthy adults (n = 16; 24.4 [20-30]) |  |  |  | After |  | NA |  |
| Roig-Hierro et al., 2023 | 32 | Healthy adults (n = 30; 22.9 ± 0.3) | Shuttle run | High<br>13 min - HIIT<br>3 min at 80% age-predicted MHR<br>2min at 60% age-predicted MHR | Motor acuity | After | 7d | NA | ↗ |
| Singh et al., 2016 | 23 | Healthy adults (n = 13; avg = 27 yrs) | Lower-limb cycling | Moderate<br>20 min - CONT<br>65-70% age-predicted MHR | Motor adaptation | Before | NA | ↔ | NA |
| Skriver et al., 2014 | 32 | Healthy adults (n = 16; 24.1 ± 3.4) | Lower-limb cycling | High<br>20 min - HIIT<br>3min at high intensity<br>2min at 50W<br>Based on measured lactate level<br>Workload (200-315) | Motor acuity | Before | 1h, 24h, 7d | ↔ | ↗ for 24h and 7d |
| Snow et al., 2016 | 16 | Healthy adults (n = 16; 25.7 ± 3.1) | Lower-limb cycling | Moderate<br>30 min - CONT<br>60% measured VO<sub>2</sub> peak | Motor acuity | Before | 24h | ↔ | ↔ |
| Statton et al., 2015 | 24 | Healthy adults (n = 8; 22.4 ± 1.1) | Running | High<br>30 min - CONT<br>65-85% age-predicted MHR (average was 84% MHR) | Motor acuity | Before | NA | ↗ | NA |
| | 20 | Healthy adults (n = 8; 24.9 ± 1.1) | Running (3 sessions) | | | | 24h | ↗ | ↔ |
| Stavrinos & Coxon 2017 | 24 | Healthy adults (n = 12; 24.8 ± 5.8) | Lower-limb cycling | High<br>20 min - HIIT<br>2min at 90% age-predicted HRR<br>3min at 50% age-predicted HRR | Motor acuity | Before | 5h | ↔ | ↗ |
| Steib et al., 2018 | 17 | Parkinson's disease (n = 17; 64.4 ± 6.2) | Lower-limb cycling | Moderate<br>30 min - CONT<br>60-70% measured MHR | Motor adaptation | Before | 24h | ↔ | ↗ |
| Stranda et al., 2019 | 26 | Healthy adults (n = 13; 25.3 ± 2.5) | Lower-limb cycling (12 sessions) | Moderate<br>20 min - CONT<br>65% age-predicted MHR | Motor acuity | Before | 7d | ↔ | ↔ |

**Table S1. continued**
| Study | Total N | Type of population (n of exercise groups; age ± SD years) | Physical exercise | Exercise parameters | Motor Learning Category | Exercise Timing to Acquisition | Retention | Exercise effect on motor learning compared to control | |
| --- | --- | --- | --- | --- | --- | --- | --- | --- | --- |
|  |  |  |  |  |  |  |  | Acquisition | Retention |
| Swarbrick et al., 2020 (no control) | 25 | Healthy adults (n = 12; 21.5 ± 2.1) | Lower-limb cycling | Low<br>19 min - LIIT<br>2 min at 8% W<sub>peak</sub><br>3 min at 12% W<sub>peak</sub> | Motor acuity | Before | 1h, 24h, 7d | ↔ | ↔ |
| | | Healthy adults (n = 13; 22.2 ± 3.1) | | High<br>19 min - HIIT<br>2 min at 60% measured W<sub>peak</sub><br>3 min at 90% measured W<sub>peak</sub> | | | | ↔ | ↔ |
| Taylor et al., 2024 (Active rest) | 23 | Healthy older adults (n = 11; 65.3 ± 6.3) | Lower-limb cycling | High<br>20 min - HIIT<br>3 min at 50% measured HRR<br>2 min at 90% measured HRR | Motor acuity | Before | 6h | ↘ | ↗ |
| Thomas et al., 2016 | 36 | Healthy adults (n = 12; 23.5 ± 2.3) | Lower-limb cycling | Low<br>17 min - LIIT<br>2 min at 25% measured W<sub>peak</sub><br>3 min at 45% measured W<sub>peak</sub> | Motor acuity | After | 24h, 7d | NA | ↗ for High at 24h<br>↗ for Low and High at 7d |
| | | Healthy adults (n = 12; 24.3 ± 2.3) | | High<br>17 min - HIIT<br>2 min at 60% measured W<sub>peak</sub><br>3 min at 90% measured W<sub>peak</sub> | | | | | |
| Thomas et al., 2017 | 40 | Healthy adults (n = 10; 24.7 ± 3.5) | Strength training | High<br>45-50 min<br>Resistance training program (4 RM and 8 RM) | Motor acuity | After | 1h, 24h | NA | ↘ at 1h<br>↗ at 1d |
| | | Healthy adults (n = 10; 25.9 ± 4.5) | Circuit training | High<br>45-50 min<br>(30% of 1 RM) | | After | | | |
| | | Healthy adults (n = 10; 25.9 ± 3.7) | Hockey training | High<br>48 min<br>5 x 7 min; 2 min rest | | After | | | |
| Thompson et al., 2023 | 45 | Stroke patients (n = 15; 68.1 ± 7.1) | Whole-body cycling | High<br>5 min - CONT<br>70-85% age-predicted MHR | Motor adaptation | After | 24h | NA | ↔ |
| | | Stroke patients (n = 15; 61.8 ± 7.5) | | High<br>15 min - HIIT<br>3 min at 70%-85% age-predicted MHR<br>2 min at < 70% age-predicted MHR | | | | | |
| Tomprowski and Pendleton, 2018 | 32 | Healthy adults (n = 11; 22.4 ± 3.0) | Dancing | Moderate<br>10 min - CONT<br>Simple Dance Dance Revolution<br>Intensity determined based on HR | Motor acuity | Before | 13min, 24h, 7d | ↔ | ↔ |
| | | Healthy adults (n = 10; 22.9 ± 2.9) | | Moderate<br>10 min - CONT<br>Complex Dance Dance Revolution<br>Intensity determined based on HR | | After | | NA | ↗ at 7d |
| Wanga et al., 2023 | 33 | Healthy adults (n = 11; 20.5 ± 2.3) | Lower-limb cycling | Low<br>23 min - HIIT<br>3min at 45% measured W<sub>peak</sub><br>2min at 25% measured W<sub>peak</sub> | Sequence | After | 24h, 48h | NA | ↗<br>(higher for Low) |
| | | Healthy adults (n = 11; 20.3 ± 2.3) | | High<br>23 min - HIIT<br>2min at 60% measured W<sub>peak</sub><br>3min at 90% measured W<sub>peak</sub> | | | | | |
| Wanner et al., 2020 | 50 | Healthy adults (n = 17; 25.3 ± 2.7) | Lower-limb cycling | Low<br>17 min - CONT<br>25W | Motor adaptation | Before | 24h | ↔ | ↔ |
| | | Healthy adults (n = 18; 25.1 ± 2.3) | | Moderate<br>17 min - HIIT<br>2min at 25% measured W<sub>peak</sub><br>3min at 45% measured W<sub>peak</sub> | | | | | |
| | | Healthy adults (n = 15; 25.7 ± 3.6) | | High<br>17 min - HIIT<br>2min at 60% measured W<sub>peak</sub><br>3min at 90% measured W<sub>peak</sub> | | | | | |
| Wanner et al., 2021 | 17 | Parkinson's disease (n = 8; 59.3 ± 5.4) | Lower-limb cycling | Moderate<br>25 min - CONT<br>60% maximal W<sub>peak</sub> | Motor adaptation | Before | 1d, 7d | ↔ | ↗ at 7d |
| Williams et al., 1985 | 90 | Healthy adults (n = 30; 20.9 ± 1.9) | Arm cranking | Low<br>5 min - CONT<br>WO maintained at 10W (30 rpm) (110-125 bpm) | Motor acuity | Before | 24h | ↔ | ↔ |
| | | Healthy adults (n = 30; 20.9 ± 1.9) | | High<br>5 min - CONT<br>WO maintained at 75W (90 rpm) (165-180 bpm) | | | | | |
| Yildirim et al., 2024 | 70 | Healthy adults (n = 15; 20.5 ± 0.7) | Running | Low<br>30 min - CONT<br>57%-63% estimated MHR | Motor acuity | Before | 24h, 7d | ↔ | ↔ |
| | | Healthy adults (n = 15; 20.1 ± 0.5) | | | | After | | ↔ | ↔ |
| | | Healthy adults (n = 15; 20.8 ± 1.5) | | Moderate<br>30 min - CONT<br>64%-76% estimated MHR | | Before | | ↔ | ↔ |
| | | Healthy adults (n = 15; 20.9 ± 1.0) | | | | After | | ↔ | ↔ |

CONT = continuous; HRR = Heart rate reserve; W = Watts; HIIT = high-intensity interval training; LIIT = light-intensity interval training; HR = Heart Rate; MHR = maximal heart rate; PPO = peak power output; rpm = revolutions per minute; bpm = beats per minute; RM = repetition maximum; Before = exercise was performed before the motor task practice; After = exercise was performed after the motor task practice; min = minutes; h = hours; d = days; NA = not available; ↗ = enhancement following exercise as reported by the authors; ↘ = decrease following exercise as reported by the authors; ↔ = no change following exercise as reported by the authors.

## Supplementary Material 2

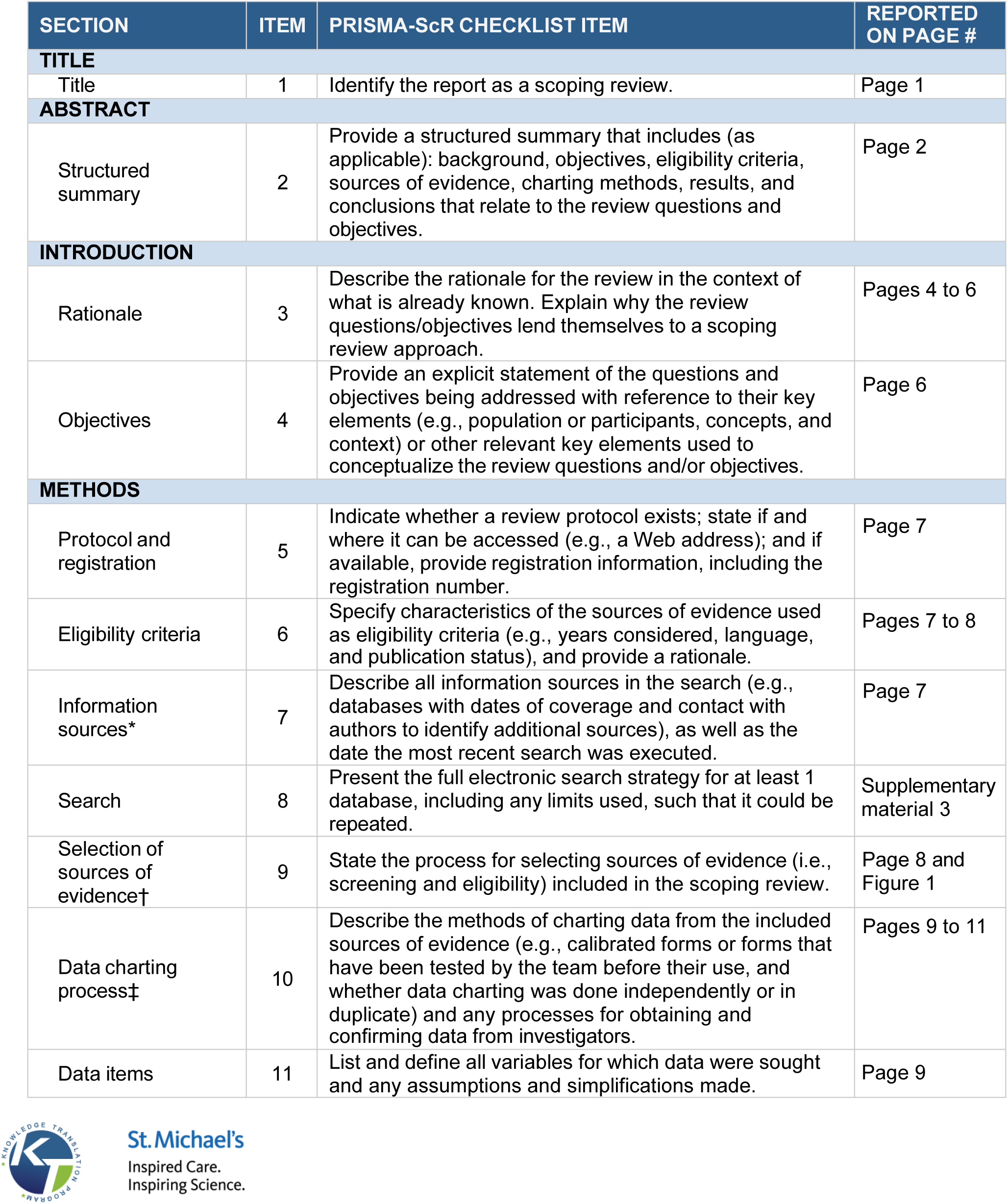
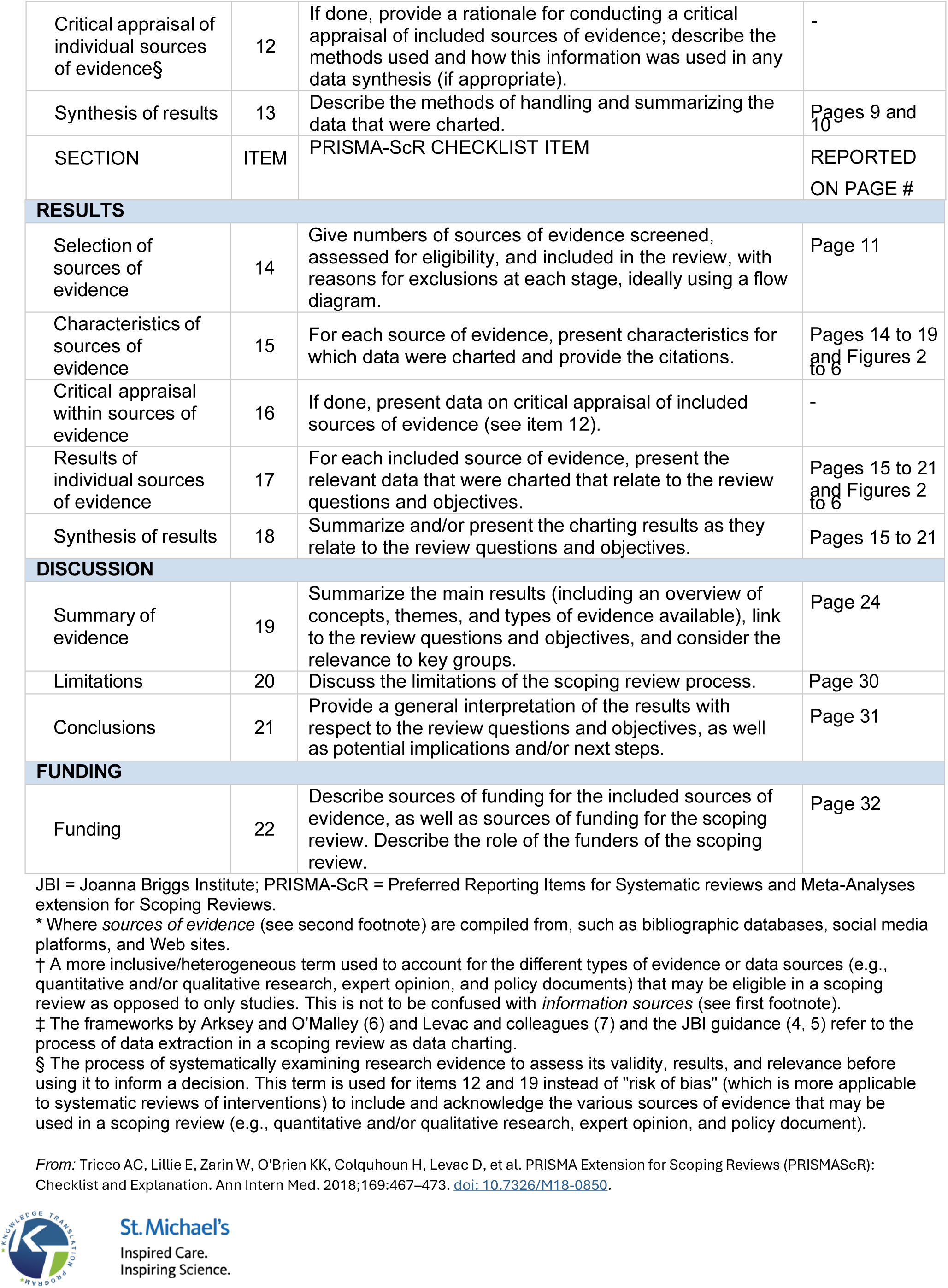

## Supplementary Material 3

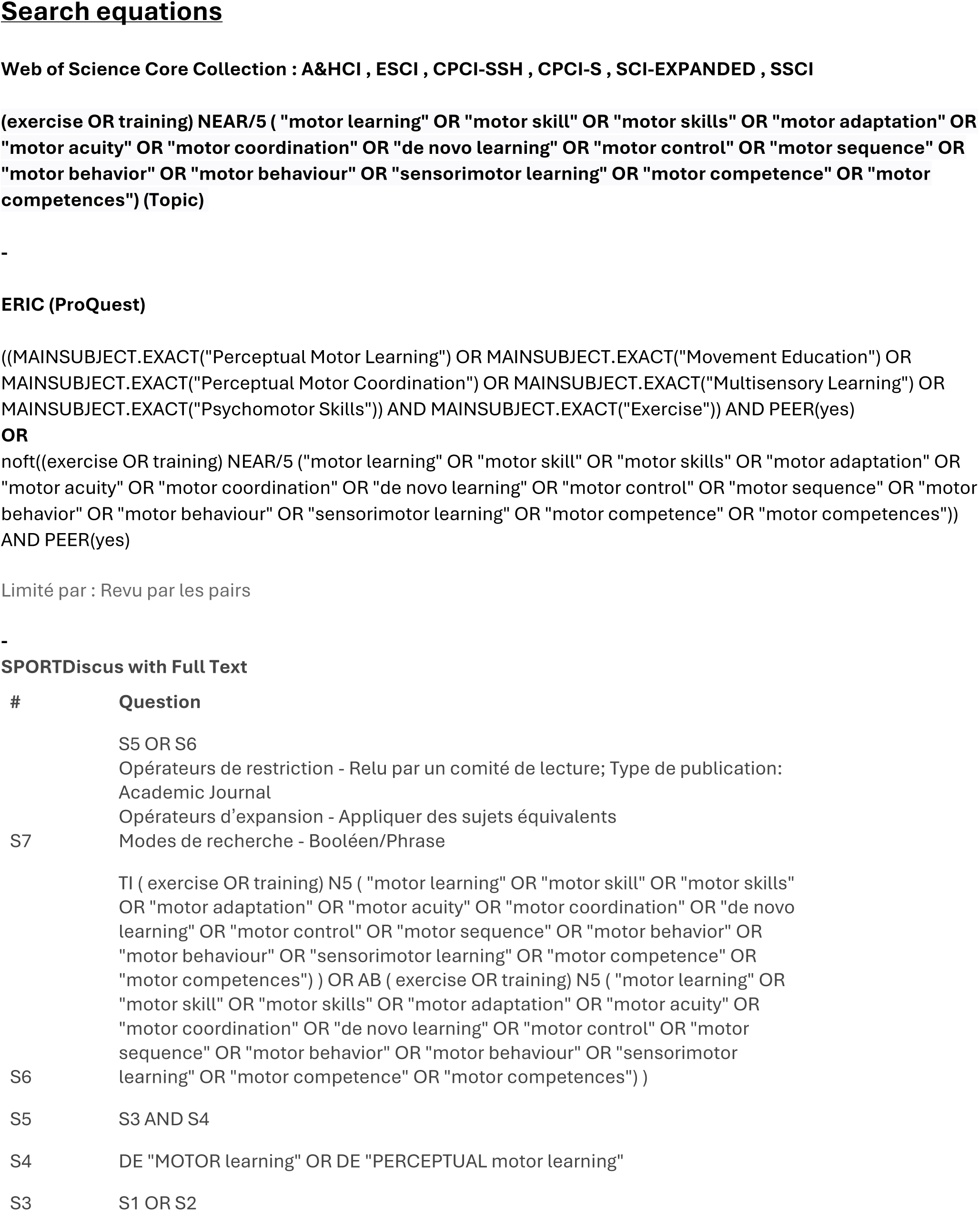
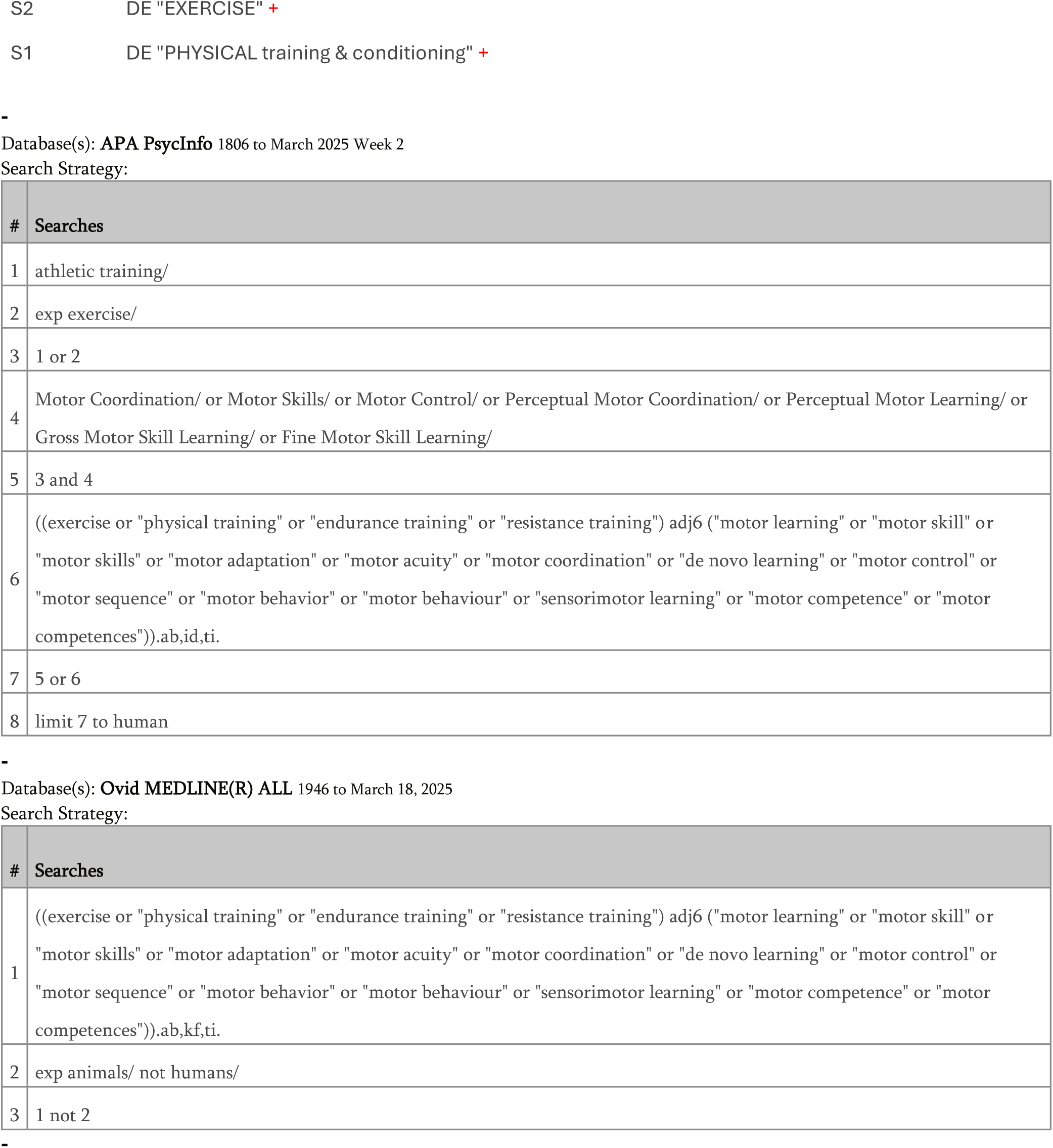
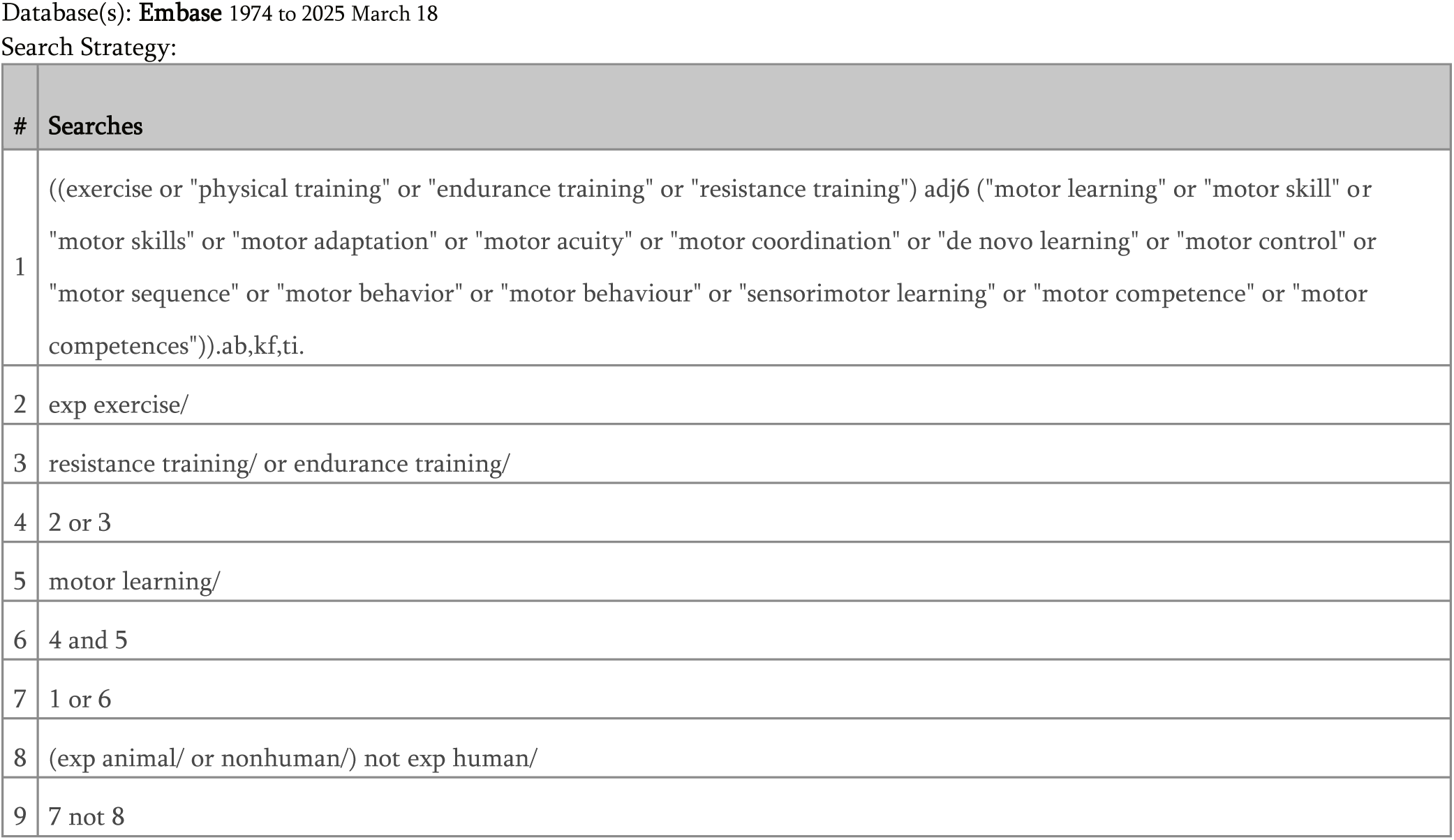

## Supplementary Material 4

Data extraction sheet is attached to this manuscript on BioRxiv

